# The monomer/dimer switch modulates the activity of plant adenosine kinase

**DOI:** 10.1101/2024.12.05.626941

**Authors:** David Jaroslav Kopečný, Armelle Vigouroux, Jakub Bělíček, Martina Kopečná, Radka Končitíková, Jaroslava Friedecká, Václav Mik, Jan František Humplík, Marine Le Berre, Stephan Plancqueel, Miroslav Strnad, Klaus von Schwartzenberg, Ondřej Novák, Solange Moréra, David Kopečný

**Affiliations:** Department of Experimental Biology, Faculty of Science, Palacký University, Olomouc CZ-78371, Czech Republic; Université Paris-Saclay, CEA, CNRS, Institute for Integrative Biology of the Cell (I2BC,) Gif-sur-Yvette F-91198, France; Laboratory of Growth Regulators, Faculty of Science, Palacký University & Institute of Experimental Botany of the Czech Academy of Sciences, Šlechtitelů 11, Olomouc CZ-78371, Czech Republic; Department of Chemical Biology, Faculty of Science, Palacký University, Olomouc CZ-78371, Czech Republic; Institute for Plant Science and Microbiology, Universität Hamburg, 22609 Hamburg, Germany

**Keywords:** Crystal structure, cytokinin, overexpression, purine, *Physcomitrella patens*, riboside, *Zea mays*

## Abstract

**Highlight:** The switch from active monomers to inactive dimers in plant ADKs impacts overall enzyme activity and represents a novel negative feedback-loop mechanism to maintain steady levels of adenosine and AMP.

Adenosine undergoes ATP-dependent phosphorylation catalyzed by adenosine kinase (ADK). In plants, ADK also phosphorylates cytokinin ribosides, transport forms of the hormone. Here, we investigated the substrate preferences, oligomeric states and structures of ADKs from moss (*Physcomitrella patens*) and maize *(Zea mays*) alongside metabolomic and phenotypic analyses. We showed that dexamethasone-inducible *ZmADK* overexpressor lines in Arabidopsis can benefit from a higher number of lateral roots and larger root areas under nitrogen starvation. We discovered that maize and moss enzymes can form dimers upon increasing protein concentration, setting them apart from the monomeric human and protozoal ADKs. Structural and kinetic analyses revealed a catalytically inactive unique dimer. Within the dimer, both active sites are mutually blocked. The activity of moss ADKs, exhibiting a higher propensity to dimerize, was tenfold lower compared to maize ADKs. Two monomeric structures in a ternary complex highlight the characteristic transition from an open to a closed state upon substrate binding. This suggests that the oligomeric state switch can modulate the activity of moss ADKs and likely other plant ADKs. Moreover, dimer association represents a novel negative feedback mechanism, helping to maintain steady levels of adenosine and AMP.

## INTRODUCTION

Cytokinins (*N^6^*-substituted adenine/adenosine derivatives) regulate cell division and many developmental events (Mok and Mok, 2001; Sakakibara, 2006) They are effective long-distance signaling molecules transported within the xylem and phloem. The xylem-transported cytokinin is predominantly trans-zeatin riboside (*t*ZR), while isopentenyl adenosine (iPR) and cis-zeatin riboside (cZR) appear in the phloem. Mainly bases (iP, *t*Z, *c*Z) bind to histidine kinase receptors and trigger the signaling cascade leading to primary hormone responses (Sakakibara, 2006). Cytokinin ribosides are thus considered transport forms of active hormone (Hirose et al., 2008) and together with other nucleosides they are transported to the cytosol by members of the equilibrative nucleoside transporter (ENT) family (Wormit et al., 2004; Girke et al., 2014). Increased nucleoside import into the cytosol has been reported upon nitrogen starvation (Cornelius et al., 2012; Melino et al., 2018).

Nucleosides are hydrolyzed by nucleosidases (NRHs) to corresponding bases and ribose (Jung et al., 2009; Jung et al., 2011; Kopečná et al. 2013; Baccolini and Witte, 2019). Purine nucleosides and bases are recycled to nucleoside monophosphates (a salvage pathway), which preserves energy instead of *de novo* synthesis (Zrenner et al., 2006; Ashihara et al., 2018). An important role has been shown for adenosine (Ado) kinase (ADK, Moffatt et al., 2000; Moffatt et al., 2002; Schoor et al., 2011) and adenine phosphoribosyltransferase (APT) (Moffatt and Somerville, 1988; Allen et al., 2002), which both produce Ado monophosphate (AMP) (Figure 1). ADK regulates intracellular Ado pools and extracellular adenylate levels. It phosphorylates Ado-derived analogs at the 5’-hydroxyl group, including cytokinin ribosides and uses ATP as a co-substrate as reported for isoforms from wheat, yellow lupin, moss, Arabidopsis or tobacco (Chen and Eckert, 1977; Guranowski, 1979; von Schwartzenberg et al., 1998; Moffatt et al., 2000; Kwade et al., 2005). Resulting cytokinin riboside monophosphates (iPRMP, *t*ZRMP, *c*ZRMP) can be hydrolyzed into an active hormone by the LONELY GUY phosphoribohydrolase (LOG) (Kuroha et al., 2009).

**Figure 1.**
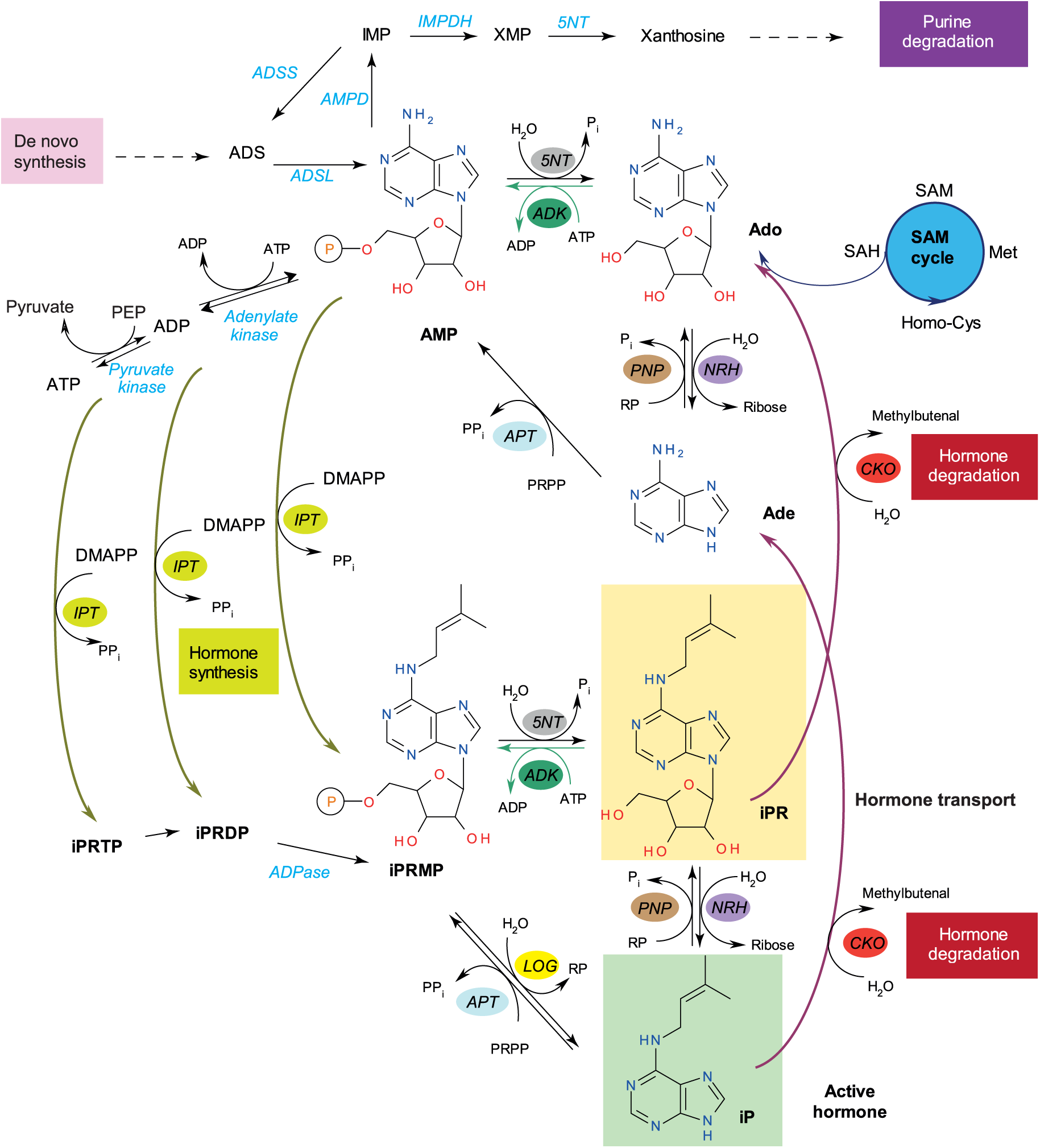
A scheme of adenosine and cytokinin riboside interconversion to their nucleobase and monophosphate derivatives. Interconversion between nucleoside monophosphate and nucleoside is catalyzed by adenosine kinase (ADK) and 5’nucleotidase (5NT). NRH and purine nucleoside phosphorylase (PNP) facilitate the conversion between bases and ribosides. Adenine phosphoribosyltransferases (APT) and cytokinin phosphoribohydrolase, known as LONELY GUY (LOG), catalyze the conversion between nucleobase and riboside monophosphate. Additional enzymes involved in the process include AMP dehydrogenase (AMPD), ADS lyase (ADSL), ADS synthetase (ADSS), cytokinin oxidase/dehydrogenase (CKO), IMP dehydrogenase (IMPDH), and isopentenyl transferase (IPT).

Maintenance of AMP levels is essential for the further interconversion to ADP and ATP, energy-rich molecules critical for all metabolic pathways. AMP is also needed to maintain the purine nucleotide pools and catabolism. Although AMP is also *de novo* synthesized in plastids, *adk* and *apt* mutants were shown to exhibit a reduced or a complete loss of fertility, changes in transmethylation reactions, and abnormal cell walls, confirming their essential role in AMP production (Allen et al., 2002; Schoor et al., 2011). ADK-deficient plants had impaired root growth, stamen and petal development, crinkled rosette leaves, reduced size of meristems and accumulated cytokinin ribosides. The primary sources of Ado, apart from the import into cells and degradation of RNA in vacuoles, are *S*-adenosyl homocysteine (SAH) hydrolase and purine nucleoside phosphorylase (PNP) (Sauter et al., 2013; Bromley et al., 2014) that catalyzes the ribosylation reaction of adenine. ADK must steadily remove Ado to prevent the feedback inhibition of SAH hydrolase, which belongs to the *S*-adenosyl methionine (SAM) cycle, and in consequence, to maintain the rate of SAM-dependent transmethylation reactions (Moffatt et al., 2002). The primary source of adenine, apart from the NRH reaction, is the hydrolysis of methyl-5’-thioAdo in reactions leading from SAM to polyamines, ethylene and nicotinamide (Sauter et al., 2013). Finally, Ado and Ade appear as products of cytokinin degradation by cytokinin oxidase/dehydrogenase (CKO/CKX, reviewed in Schmülling et al., 2003), while AMP, ADP or ATP are used by isopentenyl transferase (IPT) for cytokinin biosynthesis (Figure 1).

Ado is one of the most ancient signaling molecules. Moreover, Ado and other purine and pyrimidine nucleotides trigger signaling through purinergic receptors. Four subtypes of P1 receptors bind Ado, whereas two subtypes of P2 receptors bind tri and dinucleotides (Burnstock, 2007, Khakh and Burnstock, 2009). The first plant Ado receptor was discovered only ten years ago (Choi et al., 2014). Because plants lack Ade/Ado deaminase, only AMP is deaminated to inosine monophosphate (IMP) by AMP deaminase (Xu et al., 2005; Han et al., 2006). IMP then undergoes either oxidation to xanthosine monophosphate (XMP) and further to xanthine into the purine degradation pathway or is converted to adenylosuccinate (ADS) by ADS synthetase and further back to AMP by ADS lyase (Hatch, 1966) (Figure 1). It is known that protozoan parasites such as *Toxoplasma gondii*, which cannot synthesize purines *de novo*, access nucleosides from their mammalian host mainly via ADK, and use them for DNA synthesis. (Schwartzman and Pfefferkorn, 1982).

There are several structures of ADK from *T. gondii* (Schumacher et al., 2000; Zhang et al., 2007)*, Trypanosoma brucei* (Kuettel et al, 2011; Timm et al., 2014) or human (Mathews et al., 1998; Muchmore et al., 2006). In each case, the enzyme is composed of large and small domains. The large domain comprises the low-affinity binding co-substrate ATP site and the groove between the large and small domains contains the high-affinity Ado-binding site. Upon substrate binding, the small domain moves and seals the cavity from an open to closed conformation. Other nucleosides, such as 2-deoxyriboside and arabinoside derivatives, were very poor substrates for rabbit ADK (Miller et al., 1979a). Ado analog adenine arabinoside (AraA) was identified as an efficient inhibitor of TgADK *in vivo* (Pfefferkorn and Pfefferkorn, 1976). However, there are other more potent ligands, such as *N*^6^-(*p*-methoxybenzoyl)Ado or 7-iodo-7-deazaAdo (iodotubercidin) (Iltzsch et al., 1995).

Here, we focused on the metabolism of nucleosides through the function of ADK from the seed plant model maize (*Zea mays*) and the early divergent model plant moss *Physcomitrella patens*, *in vitro* and *in planta*. Given the known negative effects of lower ADK activity on plant growth (Allen et al., 2002; Schoor et al., 2011), we prepared homozygous dexamethasone-inducible *ZmADK* overexpressor lines (for all three maize *ADK* genes) in *Arabidopsis thaliana* to verify the hypothesis that plants should benefit from increased ADK activity upon stress conditions. Our findings demonstrated the benefits of increased ADK activity *in planta.* We showed the crucial role of the monomer for ADK activity and discovered, for the first time, the ability of plant ADKs to associate into an inactive dimer. The existence of the dimer was further confirmed by X-ray crystallography on ZmADK2 and the structure explains why this oligomeric state is not functional. Furthermore, structural analysis of two ternary complexes of ZmADK3 and PpADK1 together with kinetic and binding studies provide insight into nucleoside recognition and conformational changes of the enzyme.

## MATERIALS AND METHODS

### Cloning, expression and purification of ADKs from P. patens and Z. mays

The total RNA was extracted from various maize organs (*Zea mays* cv. Cellux, Morseva) and from Physcomitrella (‘Gransden 2004’ strain) at the protonema stage using the RNAqueous kit and plant RNA Isolation Aid. The cDNA was synthesized with Superscript III RT and oligo dT primers (www.thermofisher.com). The sequences encoding maize ADKs, specifically *ZmADK1* (Zm00001d051157, 1029 bp), *ZmADK2* (Zm00001d017271, 1029 bp) and *ZmADK3* (Zm00001d003017, 1035 bp) were cloned with Platinum™ SuperFi™ DNA Polymerase (www.thermofisher.com) into a pCDFDuet His-tag vector (Novagen, www.merckmillipore.com) using two forward primers containing *Sac*I site and a common reverse primer with *Kpn*I site (5′-ATTGAGCTCGATGGCGGCGAGCGAGGGCGT-3′ and 5′-ATTGAGCTCGGCCAGCAGCGGCTACGAAGGGA-3′; 5′-ACTGGTACCCTAGTTGAAGTCAGGCTTCT-3′). Similarly, two sequences for moss ADKs were cloned, namely *PpADK1* (Pp3c3_10800, 1032 bp) and *PpADK2* (Pp3c13_10550, 1026 bp) using primer pairs containing *Sac*I/*Kpn*I and *BamH*1/*Xho*1 sites (5′-ATAGAGCTCGATGGCGTCCGAGGGTGTGCTTTT-3′ and 5′-ACTGGTACCCTACTGACTTTCGAAGGATGGTT-3′; 5′-CATGGATCCGATGGCGTCCGAAGGAGTG-3′ and 5′-CATCTCGAGCTACTCCGACTGAAAACAAG-3′). The constructs were transformed into T7 Express *I^q^ E. coli* cells (www.neb.com).

Protein production was conducted at 200 rpm and 18°C overnight with 0.5 mM isopropylthio-β-galactoside. The enzymes were subsequently purified using a Nickel-HiTrap IMAC FF column (https://www.cytivalifesciences.com) on an NGC Medium-Pressure Liquid Chromatography System (https://www.bio-rad.com) and further purified by gel filtration chromatography on a Superdex 200 10/30 HR column (www.gelifesciences.com) into 20 mM Tris-HCl buffer, pH 7.5, 100 mM NaCl with 1% glycerol. The yield after purification was approximately 4-6 mg per 100 ml culture. Cells were analyzed for the expression of the His-tagged protein by NuPAGE (www.thermofisher.com) and using activity measurements. NuPAGE electrophoresis was performed in MES buffer on 4–12% Bis-Tris gel. Protein content was quantified using the Coomassie plus (Bradford) protein assay kit (www.thermofisher.com).

### Site-directed mutagenesis of ZmADK3

All *ZmADK3* mutants were cloned using tail-to-tail oriented phosphorylated primers, with the mutation being located at the 5′ end of one of the primers by a PCR reaction in 30 cycles. The following mutants were cloned: D19A (5′-P-GCCATCTCCGCTGTCGTCGA-3′, 5′-P-GAGGAGGGGGTTCCCCATTC-3′), R132A (5′-P-GCGTCACTTATTGCTAACTTGTCT-3′, 5′-TTCTCCACCAACAACACAAAC-3′), N222A (5′-P-GCAGAAACTGAGGCGAGGACC-3′, 5′-P-TCCAAAGATGTAATCCACGTACGG-3′), E225A (5′-P-GCGAGGACCTTTGCTAAAGT-3′, 5′-P-GCGAGTTTCATTTCCAAAGATGT-3′), N295A (5′-P-GCAGGTGCAGGTGATGCGTTC-3′, 5′-P-GGTGTCAACAAGCTTCTCCTTA-3′) and D299A (5′-P-GCGGCGTTCGTTGGTGGCTT-3′, 5′-P-ACCTGCACCATTGGTGTCA-3′). The products were treated with *Dpn*I restriction enzyme, gel purified and ligated using T4 DNA ligase (www.thermofisher.com). Clones were transformed into T7 express *I^q^ E. coli* competent cells.

### Synthesis of N^6^-methylAdo, N^6^, N^6^-dimethylAdo, N^6^-isopropylAdo and N^6^-isobutylAdo

*N^6^*-methylAdo was prepared by mixing a suspension of 6-chloropurine riboside (1 g, 3.49 mmol) heated in 33% methylamine solution in absolute ethanol (4.33 ml, 34.9 mmol) at 90 °C for 4 h. *N^6^, N^6^*-dimethylAdo was prepared by mixing a suspension of 6-chloropurine riboside (1 g, 3.49 mmol) with dimethylamine hydrochloride (0.569 g, 6.98 mmol) and triethylamine (1.7 ml, 12.21 mmol) and heating in 20 ml of methanol at 100 °C for 4 h. *N^6^*-isopropylAdo and *N^6^-*isobutylAdo were prepared in a one-step reaction by heating of 6-chloropurine riboside with the corresponding amine in *n*-propanol at 85 °C, with triethylamine serving as an auxiliary base. A suspension of 6-chloropurine riboside (100 mg, 0.349 mmol), either isopropylamine (5 eq. 149 μl, 1.745 mmol) or isobutylamine (1.2 eq., 42 µl, 0.419 mmol), along with triethylamine (1.2 eq., 121 μl, 0.872 mmol), was heated in pressure tubes in 0.25 M *n-*propanol at 85 °C for four h.

The reaction mixtures were concentrated under reduced pressure and the product was purified by silica column chromatography using chloroform/methanol as a mobile phase, starting from 25:1 (v/v) with methanol gradient. The reaction progress was monitored by thin-layer chromatography using Silica F254 aluminium plates (www.avantorsciences.com), and the spots were detected by 254 nm UV light and/or by vanillin staining. Chromatographic purity and mass of the prepared compounds were determined using the AQUITY UPLC H-Glass system (www.waters.com) on a Symmetry C18 column (150 × 2.1 mm, particle size 5 µ, www.waters.com). NMR spectra were recorded on an ECA-500 (www.jeol.com) operating at a frequency of 500 MHz (^1^H) and 125 MHz (^13^C). Substances were dissolved in DMSO-*d*_6_ and chemical shifts were calibrated to the residual/solvent peak DMSO-*d*_5_ −2.49 ppm for ^1^H and DMSO-*d*_6_ - 39.5 ppm for ^13^C. High-resolution mass spectra were obtained on 1290 Infinity II Autoscale preparative LC/MSD System (www.agilent.com) coupled simultaneously to 6230 Time-of-Flight LC/MS mass spectrometer (LC/TOF, Agilent) with an electrospray ionization source Dual AJS ESI.

Final characteristics of *N^6^-*methylAdo (6-methylamino-9*H-*purine-9-ß-D-riboside): white solid, yield: 942 mg (96%), HPLC-UV/VIS retention time, purity (min., %): 4.73, 99.9. ESI^+^-MS *m/z* (rel. int. %, ion): 282.4 (100, [M+H]^+^). ^1^H-NMR (500 MHz, DMSO-*d*_6_) δ (ppm): 2.94 (s, 3H,-NHC<u>H</u>_3_), 3.54 (ddd, *J* = 12.2, 7.3, 3.7 Hz, 1H, rib H5‘), 3.66 (dt, *J* = 12.0, 3.8 Hz, 1H, rib H5‘), 3.95 (q, *J* = 3.3 Hz, 1H, rib H4‘), 4.13 (td, *J* = 4.6, 3.1 Hz, 1H, rib H3‘), 4.59 (q, *J* = 5.9 Hz, 1H, rib H2‘), 5.18 (d, *J* = 4.3 Hz, 1H, rib C3‘O<u>H</u>), 5.42-5.44 (m, 2H, rib C2‘ O<u>H</u>, rib C5‘ O<u>H</u>), 5.87 (d, *J* = 6.7 Hz, 1H, rib H1‘), 7.83 (bs, 1H,-N<u>H</u>CH_3_), 8.22 (s, 1H, pur H2), 8.33 (s, 1H, pur H8). ^13^C-NMR (125 MHz, DMSO-*d*_6_) δ (ppm): 27.0 (-NH<u>C</u>H_3_), 61.7 (rib C5‘), 70.7 (rib C3‘), 73.5 (rib C2‘), 85.9 (rib C4‘), 87.9 (rib C2‘), 119.9 (pur C5), 139.7 (pur C8), 148.0 (pur C4), 152.5 (pur C2), 155.1 (pur C6). HRMS (ESI/TOF) m/z: [M + H]^+^ Calcd for C_11_H_16_N_5_O_4_ 282.1197; Found 282.1199.

Final characteristics of *N^6^, N^6^-*dimethylAdo (6-(dimethylamino)-9*H-*purine-9-ß-D-riboside): white solid, yield: 995 mg (97%), HPLC-UV/VIS retention time, purity (min., %): 6.44, 99.9. ESI^+^-MS *m/z* (rel. int. %, ion): 296.4 (100, [M+H]^+^). ^1^H-NMR (500 MHz, DMSO-*d*_6_) δ (ppm): 3.44 (vbs, 6H,-NH(C<u>H</u>_3_)_2_), 3.54 (ddd, *J* = 12.0, 7.0, 3.7 Hz, 1H, rib H5‘), 3.66 (dt, *J* = 12.1, 4.1 Hz, 1H, rib H5‘), 3.95 (q, *J* = 3.4 Hz, 1H, rib H4‘), 4.13 (td, *J* = 4.7, 3.4 Hz, 1H, rib H3‘), 4.56 (q, *J* = 5.7 Hz, 1H, rib H2‘), 5.17 (d, *J* = 4.6 Hz, 1H, rib C3‘ O<u>H</u>), 5.36 (dd, *J* = 6.7, 4.6 Hz, 1H, rib C5‘ O<u>H</u>), 5.43 (d, *J* = 6.4 Hz, 1H, rib C2‘ O<u>H</u>), 5.90 (d, *J* = 6.1 Hz, 1H, rib H1‘), 8.20 (s, 1H, pur H2), 8.36 (s, 1H, pur H8). ^13^C-NMR (125 MHz, DMSO-*d*_6_) δ (ppm): 37.9 (2× C,-N(<u>C</u>H_3_)_2_), 61.5 (rib C5‘), 70.5 (rib C3‘), 73.5 (rib C2‘), 85.7 (rib C4‘), 87.8 (rib C1‘), 119.8 (pur C5), 138.6 (pur C8), 149.9 (pur C4), 151.7 (pur C2), 154.3 (pur C6). HRMS (ESI/TOF) m/z: [M + H]^+^ Calcd for C_12_H_18_N_5_O_4_ 296.1353; Found 296.1351.

Final characteristics of *N^6^-*isopropylAdo (6-isopropylamino-9*H*-purine-9-ß-D-riboside): white solid, yield of 93 mg (86%), HPLC-UV/VIS retention time, purity (min., %): 7.41, 97.3. ESI^+^-MS *m/z* (rel. int. %, ion): 310.3 (100, [M+H]^+^). ^1^H-NMR (500 MHz, DMSO-*d*_6_) δ (ppm): 1.20 (d, *J* = 6.7 Hz, 6H,-NHCH(C<u>H</u>_3_)_2_), 3.54 (ddd, *J* = 12.1, 7.5, 3.7 Hz, 1H, rib H5‘), 3.66 (dt, *J* = 12.1, 4.1 Hz, 1H, rib H5‘), 3.95 (q, *J* = 3.3 Hz, 1H, rib H4‘), 4.13 (td, *J* = 4.8, 3.0 Hz, 1H, rib H3‘), 4.43 (bs, 1H, - NHC<u>H</u>(CH_3_)_2_), 4.60 (td, *J* = 6.3, 5.0 Hz, 1H, rib H2‘), 5.19 (d, *J* = 4.6 Hz, 1H, rib C3‘ O<u>H</u>), 5.43-5.46 (m, 2H, rib C2‘ O<u>H</u>, rib C5‘ O<u>H</u>), 5.86 (d, *J* = 6.1 Hz, 1H, rib H1‘), 7.66 (bd, *J* = 4.9 Hz, 1H, - N<u>H</u>CH(CH_3_)_2_), 8.19 (bs, 1H, pur H2), 8.33 (s, 1H, pur H8). ^13^C-NMR (125 MHz, DMSO-*d*_6_) δ (ppm): 22.3 (2× C,-NHCH(<u>C</u>H_3_)_2_), 41.3 (-NH<u>C</u>H(CH_3_)_2_), 61.7 (rib C5‘), 70.7 (rib C3‘), 73.4 (rib C2‘), 85.9 (rib C4‘), 87.9 (rib C1‘), 119.7 (pur C5), 139.6 (pur C8), 148.3 (pur C4), 152.4 (pur C2), 153.9 (pur C6). HRMS (ESI/TOF) m/z: [M + H]^+^ Calcd for C_13_H_20_N_5_O_4_ 310.1510; Found 310.1525.

Final characteristics of *N^6^-*isobutylAdo (6-isobutylamino-9*H*-purine-9-ß-D-riboside): white solid, yield: 80 mg (71 %); HPLC-UV/VIS retention time, purity (min., %): 9.09, 99.9. ESI^+^-MS *m/z* (rel. int. %, ion): 324.4 (100, [M+H]^+^). ^1^H-NMR (500 MHz, DMSO-*d*_6_) δ (ppm): 0.87 (d, *J* = 6.9 Hz, 6H,-NHCH_2_CH(C<u>H</u>_3_)_2_), 1.92-1.98 (m, 1H,-NHCH_2_C<u>H</u>(CH_3_)_2_), 3.28 (bt, *J* = 5.3 Hz, 2H, - N<u>H</u>CH_2_CH(CH_3_)_2_), 3.54 (ddd, *J* = 12.2, 7.4, 3.7 Hz, 1H, rib H5‘), 3.66 (dt, *J* = 12.1, 3.9 Hz, 1H, rib H5‘), 3.95 (q, *J* = 3.2 Hz, 1H, rib H4‘), 4.13 (td, *J* = 4.5, 3.2 Hz, 1H, rib H3‘), 4.59-4.62 (m, 1H, rib H2‘), 5.18 (d, J = 4.6 Hz, 1H, rib C3‘ O<u>H</u>), 5.44 (bd, *J* = 6.4 Hz, 2H, rib C2‘ O<u>H</u>, rib C5‘ O<u>H</u>), 5.86 (d, *J* = 6.4 Hz, 1H, rib H1‘), 7.95 (bt, *J* = 5.3 Hz, 1H,-N<u>H</u>CH_2_CH(CH_3_)_2_), 8.18 (bs, 1H, pur H2), 8.33 (s, 1H, pur H8). ^13^C-NMR (125 MHz, DMSO-*d*_6_) δ (ppm): 20.1 (2× C,-NHCH_2_CH(<u>C</u>H_3_)_2_), 27.8 (-NHCH_2_<u>C</u>H(CH_3_)_2_), 47.2 (-NH<u>C</u>H_2_CH(CH_3_)_2_), 61.7 (rib C5‘), 70.7 (rib C3‘), 73.4 (rib C2‘), 86.0 (rib C4‘), 88.0 (rib C1‘), 119.7 (pur C5), 139.7 (pur C8), 148.2 (pur C4), 152.3 (pur C2), 154.8 (pur C6). HRMS (ESI/TOF) m/z: [M + H]^+^ Calcd for C_14_H_22_N_5_O_4_ 324.1666; Found 324.1665.

### Enzyme kinetics

Substrate analogs were purchased from Sigma-Aldrich (www.sigmaaldrich.com). The ADK activity was measured spectrophotometrically on an Agilent 8453 UV-Vis spectrophotometer (www.agilent.com) at 30 °C using a coupled reaction with pyruvate kinase (PK) and lactate dehydrogenase (LDH) (www.sigmaaldrich.com) (Blondin et al., 1994). In this assay, PK converts ADP produced by ADK and phosphoenolpyruvate to pyruvate. Subsequently, LDH then converts pyruvate to lactate using NADH coenzyme, with the conversion monitored at 340 nm. The reaction was conducted in 50 mM Tris-HCl buffer at pH 7.5, containing 50 mM KCl, 6 mM MgCl_2_, 300 µM PEP, 200 µM NADH, 10 mM ATP, 2 units of PK and 2.5 units of LDH, using various nucleoside substrates. Kinetic constants were determined using GraphPad Prism 8.0 software (www.graphpad.com). Each measurement was performed six times using repetitive purified enzymes. The activities of all five ADKs were also measured by HPLC (Moffatt et al., 1991). The reaction mixture comprised 50 mM Tris-HCl pH 7.5, 5 mM MgCl_2_, 0.5 mM DTT, 30 mM NaF, 5 mM ATP, 40 µM substrate and 2-150 ng of purified ADK, depending on the isoform. The reaction was incubated at 30 °C for up to 1 hour. A conversion of Ado to AMP was analyzed in UV-mode using a diode array detector after separation on a LiChrosphere 60 RP-Select B (5 µm) LiChroCART column 250-4 (www.merckmillipore.com) equilibrated with 10 mM ammonium formate at pH 3.7 and gradually eluted with 100% acetonitrile. Similarly, the conversion of cytokinin ribosides to monophosphates was followed on LiChrosphere 60 RP-Select B (5 µm) LiChroCART column 125-4 column equilibrated with 10 mM triethylamine at pH 5.4 plus 10% methanol and stepwise eluted with 100% methanol.

### Affinity, thermal stability, SAXS, AUC and DLS measurements

The MST method was used to determine the binding affinity of various ribosides to ZmADK2 and PpADK1. Proteins were fluorescently labeled with RED-tris-NTA dye (www.nanotemper-technologies.com) using a 1:1 dye/protein molar ratio. The labeled protein was adjusted to 100-300 nM in 50 mM HEPES buffer pH 7.5, 1 mM MgCl_2_ and 0.2% Tween. Measurements were performed in premium capillaries on a Monolith NT.115 instrument at 30°C with 5 sec/30 sec/5 sec laser off/on/off times, respectively, with continuous sample fluorescence recording. Thermal stability was measured by nano-differential scanning fluorimetry on a Prometheus NT.48 instrument (www.nanotemper-technologies.com) in various buffers covering a pH (7.0-9.5) and temperature range (25 to 95°C), and with a heating rate of 1 °C min^-1^ and using NT melting control software. Protein unfolding was measured by detecting the temperature-induced change in tryptophan fluorescence intensity at emission wavelengths of 330 and 350 ± 5 nm. The melting temperature (*T*_m_) was deduced from the maximum of the first derivative of the fluorescence ratios F350/F330.

SAXS data were measured on a SWING beamline at the SOLEIL synchrotron (www.synchrotron-soleil.fr) using an EigerX-4M detector. Exposure time was 500 ms, detector distance was 1790 mm, X-ray wavelength was 1.033 Å. Purified proteins were analyzed in 50 mM Tris-HCl, pH 7.5 at three concentrations up to 5 mg/ml. Data were analyzed using the ATSAS v3.2.1 package, namely Primus software (Manalastas-Cantos et al., 2021).

The analytical ultracentrifugation (AUC) in sedimentation velocity mode was performed using ProteomeLab XL-I analytical ultracentrifuge (Beckman Coulter, Indianapolis, IN, USA) equipped with an An-60 Ti rotor. Samples of ZmADK2 and PpADK1 were diluted in 20 mM Tris-HCl pH 7.5 with 150 mM NaCl, 10 mM MgCl_2_, 10 µM DTT and 1% glycerol and equilibrated at 4 °C overnight. Absorbance data were collected at 25 °C and at a rotor speed of 48000 rpm. Scans were collected at 280 nm in 5-min intervals and 0.003 cm spatial resolution in continuous scan mode. The partial specific volume of the protein and the solvent density and viscosity were calculated from the amino acid sequence and buffer composition, respectively, using the Sednterp software (http://bitcwiki.sr.unh.edu). The data were analyzed with the continuous c(s) distribution model implemented in the program SEDFIT 15.01b, using a confidence level of 0.95 for the regularization procedure (Schuck, 2000). For, was used. The plots of c(s) distributions were created in GUSSI 1.3.1 (Brautigam, 2015).

The hydrodynamic diameter was determined by DLS in 20 mM Tris-HCl pH 7.5 and 1 mM MgCl_2_ alone and in the presence of 2 mM ATP or 2 mM AP5A at 22 °C. Measurement was performed using the Zetasizer Nano ZS (Malvern Instruments, UK). Hydrodynamic diameter values were calculated by Zetasizer Software v7.13 (173° angle measurement, approximation fit to the sphere).

### Crystallization and structure determination

Crystallization conditions of all three maize ADKs were screened using Qiagen kits (www.qiagen.com) and Morpheus I screen (https://moleculardimensions.com) with a Cartesian nanodrop robot (Genomic solutions) or a Mosquito (SPT Labtech). Crystals of ZmADK2 were obtained in hanging drops by mixing equal volumes of protein solution (9 mg ml^-1^) and a precipitant solution containing 2.1M ammonium sulfate, 0.2M NaF and 0.1M sodium acetate at pH 5.5. Crystals were soaked with 10 mM AMP-PCP for 10 minutes. ZmADK3 (11 mg ml^-1^) was cocrystallized with 10 mM AP5A in a precipitant solution containing 20% PEG8000, 0.5M LiCl in 0.1M Tris-HCl buffer at pH 8.0. PpADK1 (23.6 mg ml^-1^) was cocrystallized in presence of 5 mM Ado and 5 mM ADP in a precipitant solution containing 37.5% mixture of MPD, PEG1000 and PEG3350 and 0.1 M carboxylic acid in 0.1M MES/Imidazole buffer pH 6.5. Crystals were transferred to a cryoprotectant solution composed of the mother liquor supplemented with 30 % glycerol (ZmADK2) or 20 % PEG400 (ZmADK3 and PpADK1) and flash-frozen in liquid nitrogen.

Diffraction data were collected at 100 K on the PROXIMA 1 and 2 beamlines at the SOLEIL synchrotron (www.synchrotron-soleil.fr). Intensities were integrated using the XDS program (Kabsch, 2010) and further reprocessed by Staraniso (Tickle et al., 2016). Data quality was assessed using the correlation coefficient *CC*_1/2_ (Karplus and Diederichs, 2012) (Table S1). Crystal structures were determined by performing molecular replacement with Phaser (Storoni et al., 2004) using the structure of human ADK (PDB 1BX4, Mathews et al., 1998; PDB 2I6B, Muchmore et al., 2006) as a search model. Models were refined with NCS restraints and TLS using Buster 2.10 (Bricogne et al., 2011) and with ligand occupancies set to 1. Electron density maps were evaluated using COOT (Emsley and Cowtan, 2004). MolProbity was used for structure validation (Chen et al., 2010). Molecular graphics images were generated using PYMOL v 2.5 (www.pymol.org). Ligand interactions were analyzed using Discovery Studio Visualizer (BIOVIA, San Diego, USA).

### Docking of cytokinin riboside into the active site of plant ADKs and human ADK

In silico docking was performed to compare the binding of Ado and iPR to the active site of plant ADK in open (ZmADK2) and closed conformation (ZmADK3, PpADK1) with that of human ADK (PDB 1BX4) in closed conformation, using FLARE v 8.0 (Cheeseright et al., 2006; Bauer and Mackey, 2019; http://www.cresset-group.com/flare/). The proteins were prepared for docking using rule-based protonation predicted for pH 7.0 and intelligent capping. Energy grids for docking were 12 × 12 × 12 Å in dimension and centered on ribose moiety of Ado ligand. Docking calculations were carried out by the Lead Finder docking algorithm, with three independent docking runs and keeping the best poses overall (Kuhn et al., 2020). The resulting ligand orientations and conformations were scored based on their binding free energies and the Lead Finder rank score (Stroganov et al., 2008).

### Quantitative PCR (qPCR) analysis

The total RNA was extracted from 3 to 13-day-old and 3-month-old maize plants (*Zea mays* cv. Celux, Morseva) using the phenol-chloroform method with TRIzol^TM^ reagent (www.thermofisher.com). Moss *Physcomitrium patens* was grown in liquid Knop’s medium and then exposed to 10 µM hormone (abscisic acid, cytokinin or synthetic auxin), 200_mM NaCl or 400_mM sorbitol for 4_days. RNA was extracted using the citrate/citric acid method. First-strand cDNA was synthesized using the LunaScript RT SuperMix Kit (www.neb.com). RNA from four biological replicates was transcribed in two independent reactions, and PCR was performed in triplicate. Diluted cDNA samples served as templates for qPCR conducted with the RT Luna Universal Probe qPCR Master Mix (www.neb.com) on a Quant Studio 5 Real-Time PCR System (www.thermofisher.com). Primers and FAM-TAM probes were designed using Primer Express 3.0 software and are listed in Table S2. Plasmids carrying *ZmADK* genes were used as a template for the calibration curve to determine the PCR efficiencies of designed probes and primer pairs and to verify their specificity. Cycle threshold values were normalized to maize elongation factor 1α and β-actin genes and amplification efficiency (Končitíková et al., 2015).

### Preparation of ZmADK-overexpressing lines in A. thaliana

ORFs of three *ZmADK*s (*Zea mays* cv. Cellux 225) were cloned into the pENTR2B cloning vector (www.thermofisher.com) using *SalI* and *XhoI* restriction sites and then subcloned into pOpON2.1 vector possessing dexamethasone-inducible expression *in planta* and derived from the pOpOff2 vector (Wielopolska et al., 2005) using Gateway LR cloning (vector/insert ratio 1/1). These constructs were introduced into *Agrobacterium tumefaciens* strain GV3101 by electroporation and then to *Arabidopsis thaliana* (Col-0 ecotype) using a floral dip method. In the T0 generation, resistant plants were selected on MS medium with kanamycin. Resistant plants were genotyped with specific primers for *ZmADKs* and *actin 8* gene (AT1G49240) as an internal control. At least twenty transgenics for each ZmADK gene were grown and lines with multiple insertions or silenced transgenics were discarded in the next T1 generation. In the T2 generation, homozygous lines were selected based on segregation. Finally, at least three T3-independent homozygous and mono-locus transformant lines harboring *pOpOn2.1:*:*ZmADK* construct were selected for each *ZmADK* gene for phenotype and metabolite analyses. The promoter function was tested by GUS staining upon immersion of 7-day-old seedlings (WT and transgenic plants) in half MS medium supplemented with 10 mM dexamethasone. The solution was removed after 72 hours and plants were GUS stained using a β-glucuronidase reporter gene staining kit (www.sigmaaldrich.com). Positive GUS staining served as validation of the functional transgenic line.

### Western blot analysis

Plant samples were frozen in liquid nitrogen and homogenized using an oscillating mill Retsch MM 400 (www.retsch.com) for 2 minutes. Samples were boiled with 2x Laemmli buffer for 10 minutes at 100 °C. NuPAGE electrophoresis was performed in MES buffer on 4–12% Bis-Tris gel in the presence of a NuPAGE reducing agent (www.thermofisher.com). Proteins were electrotransferred to PVDF membrane (www.merckmillipore.com) under semidry conditions on a Trans-Blot Turbo system (www.bio-rad.com) and using Trans-Blot Turbo RTA mini PVDF transfer kit. The membrane was washed in a blocking buffer (2% PVP-40, 0.1% (w/v) Tween 20 in PBS) and incubated with 1000 times diluted 6xHis-tag monoclonal antibody (HIS.H8, www.thermofisher.com) for one hour. Membranes were further reacted with mouse IgGκ BP-HRP conjugate (www.scbt.com) (diluted 1:2000) and visualized with Clarity Max ECL Western Blotting Substrates (BioRad) staining solution.

### Determination of purine and cytokinin metabolites

Two-week-old *ZmADK A. thaliana* overexpressors were induced by 20 µM dexamethasone and, at 72 hours after the treatment, samples were collected and purified in triplicate (10 mg FW per sample). For quantification of the purine bases, ribosides and monophosphates, the samples (20 mg FW per sample) were homogenized, extracted in cold water with 25% ammonia (ratio 4:1), and purified by solid-phase extraction using Oasis MAX column (30 mg/1 ml) (www.waters.com) with the addition of the stable-labeled internal standards (Kopečná et al., 2013). All samples were analyzed using a liquid chromatograph SciexExionLC system coupled to tandem mass spectrometer QTRAP6500+ (www.sciex.com) (LC−MS/MS) using aminopropyl column (Luna 3μm NH_2_, 100×2 mm) (www.phenomenex.com) according to a previous protocol (Karlíková et al., 2016). Quantification of cytokinin metabolites was performed according to the described method (Svačinová et al., 2012), including modifications (Plačková et al., 2017). Samples (10 mg FW per sample) were homogenized and extracted in 1 ml of modified Bieleski buffer (60% MeOH, 10% HCOOH and 30% H_2_O) together with a cocktail of stable isotope-labeled internal standards (0.2 pmol of CK bases, ribosides, *N*-glucosides, and 0.5 pmol of CK *O*-glucosides, nucleotides per sample added). The extracts were purified by an Oasis MCX column (30 mg/1 ml) (www.waters.com) and then cytokinin levels were determined by LC-MS/MS using stable isotope-labelled internal standards as a reference. Separation was performed on an Acquity UPLC i-Class System equipped with an Acquity UPLC BEH Shield RP18 column (150×2.1 mm, 1.7 μm), and the effluent was introduced into the electrospray ion source of a triple quadrupole mass spectrometer Xevo™ TQ-S MS (www.waters.com).

### Extraction and UHPLC-MS/MS quantification of selected amino acids

Plants were extracted using a previously described method (Supíková et al., 2022) with modifications. A material (10 mg of dry weight) was extracted with methanol (1 ml) containing 0.1% formic acid. The extracts were then evaporated under nitrogen, redissolved in 100 μL of 20% methanol and analyzed by ultra-high performance liquid chromatography (UHPLC-MS/MS, www.waters.com) with a PDA detector (Acquity Ultra Performance) coupled with a tandem mass spectrometer Synapt G2-Si (www.waters.com) comprising an electrospray and a quadrupole time-of-flight (QqTOF) mass analyzer. Extracts were injected on an ARION Polar C18 column (5 μm, 250 mm × 4.6 mm, www.chromservis.eu) at 30 °C. Mobile phases A (acetonitrile) and B (5 mM formic acid) were mixed in a gradient: 0 min 5% A, 0.1 min 5% A, 7 min 10% A, 12 min 35% A, 17 min 70% A, 17.5 min 100% A, 19 min 100% A, 19.5 min 5% A, 22 min 5% A. The data were acquired in a data-dependent acquisition mode. Metabolite integration and quantification with external calibration was performed using TargetLynx (www.waters.com). All standards were purchased from Sigma-Aldrich (www.sigmaaldrich.com).

### Phenotyping of root and shoot growth

To study the root phenotype of transgenics, seeds of four independent lines and WT were germinated on half MS medium. Stratified 3-day-old seedlings were transferred onto square Petri dishes containing 20 µM dexamethasone and grown for the next 11 days on the MS medium supplemented with 0.8% gellan gum without ammonium nitrate (No N) and standard (Normal N). Plants were grown at 21 °C, 70% relative humidity and light intensity of 110 µmol m^-2^ s^-1^ with a photoperiod of 16/8 hours. Length of the primary root, total root area and number of lateral roots were monitored daily with an RGB camera on vertical plates. Growth parameters were calculated. Rosette area was followed using the 24-well plates. At least 24 seedlings (biological replicates) per line and growth condition were included in each experiment. In the case of WT, 48 seedlings were analyzed. Image analysis was performed using FIJI (Schindelin et al., 2012), resulting values were statistically analyzed using t-test with Statistica software (StatSoft, USA).

## RESULTS

### Maize are monomeric while moss ADKs are dimeric in solution

Maize and moss genomes (https://phytozome-next.jgi.doe.gov/) indicate each the existence of three putative *ADK* genes. Maize *ADK* genes are located on chromosomes 4 (Zm00001d051157), 5 (Zm00001d017271) and 2 (Zm00001d003017) and are composed of 13 exons (Figure 2A).We cloned the three maize genes to produce the recombinant proteins in bacteria. ZmADK1 (342 aa) shares a sequence identity of 96.5% with ZmADK2 (342 aa) and 87.2% with ZmADK3 (344 aa). Two genes of moss *ADKs*, each composed of 13 exons, are located on chromosomes 3 (Pp3c3_10800) and 13 (Pp3c13_10550), while the third one is placed in chromosome 8 (Pp3c8_25260) and comprises 11 exons. We cloned *PpADK1* and *PpADK2* and expressed them in *E. coli*, whereas *PpADK3* cloning was unsuccessful. PpADK1 (343 aa) and PpADK2 (341 aa) display a sequence identity below 70%. Although ADK activity in moss and PpADK1 sequence have been reported in the past (von Schwartzenberg et al., 1998), there has been no in-depth enzyme characterization so far.

**Figure 2.**
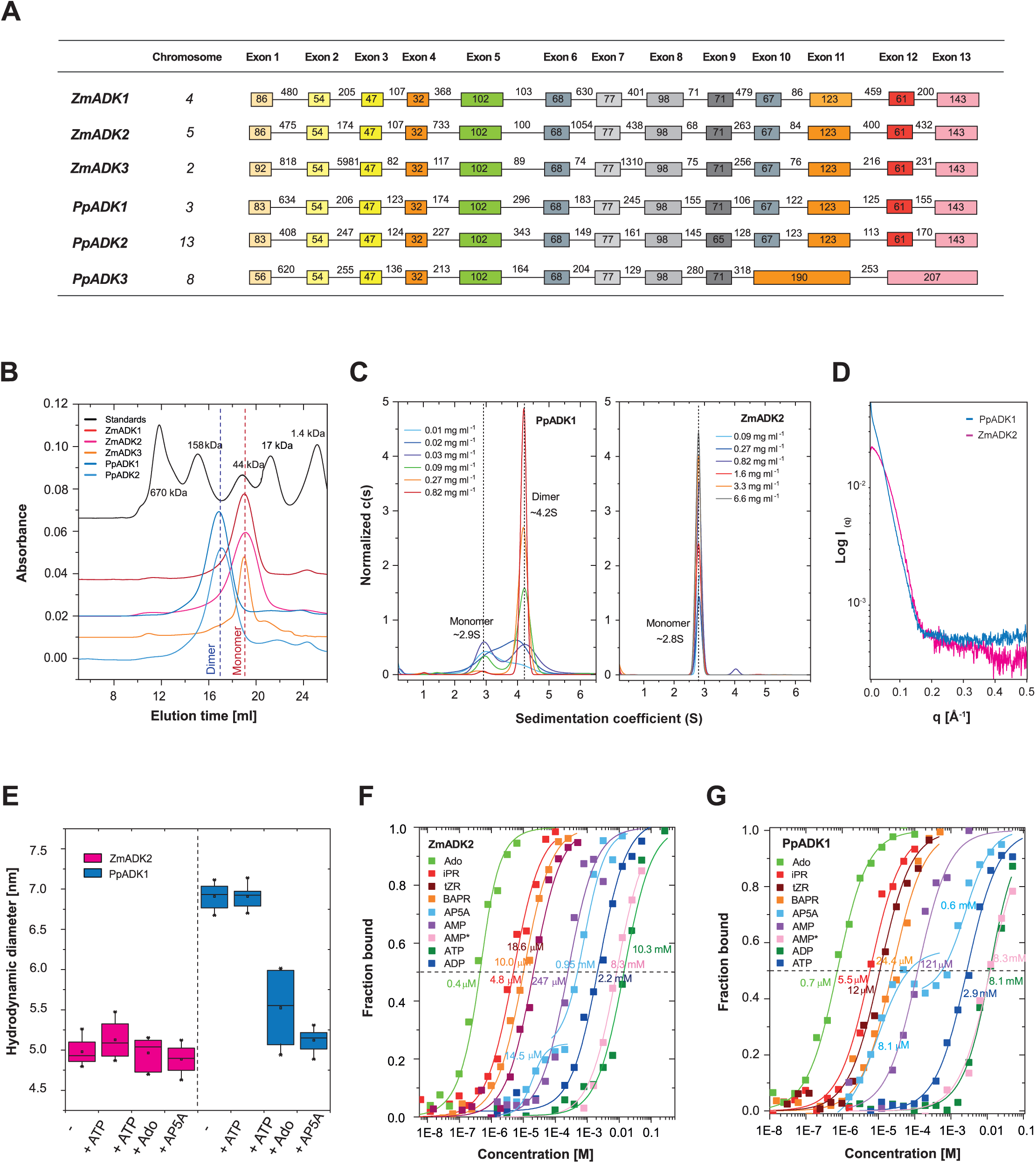
Molecular and binding properties of plant ADKs. **(A)** An overview of *ADK* gene models based on cloning in this work and genomic DNA sequences available at Phytozome (https://phytozome-next.jgi.doe.gov/). **(B)** Gel permeation chromatography profiles of five plant ADKs on a Superdex 200 10/30 HR column in 20 mM Tris-HCl buffer, pH 7.5, 100 mM NaCl, with calibration performed using a gel filtration standard (Bio-Rad). **(C)** The analytical ultracentrifugation plots of c(s) distributions for ZmADK2 and PpADK1 as analyzed in sedimentation velocity mode. Performed in 20 mM Tris-HCl pH 7.5 with 150 mM NaCl, 10 mM MgCl_2_, 10 µM DTT and 1% glycerol. Scans were collected at a rotor speed of 48000 rpm at 280 nm in 5-min intervals and 0.003 cm spatial resolution in continuous scan mode. **(D)** Plots of SAXS data measured with selected ADKs. Buffer values were subtracted. **(E)** Hydrodynamic diameter determined by DLS for selected ADKs. Measured for apoforms in 20 mM Tris-HCl pH 7.5 and 1 mM MgCl_2_ with 2 mM ATP or 2 mM AP5A at 22 °C. **(F, G)** Binding affinity curves of ZmADK2 and PpADK1 for selected riboside ligands, ADP, ATP and AP5A. Data were measured by MST in 50 mM HEPES buffer pH 7.5, 1 mM MgCl_2_ and 0.2% Tween.

We purified the five above N-terminal His-tagged ADK isoforms (theoretical MW of about 39 kDa) and investigated their molecular properties. Gel permeation chromatography analysis showed that all three maize ADKs appeared monomeric (MW of 38 ± 3 kDa) (Figure 2B) as well as dimeric in small proportion at higher concentrations (Figure S1). In contrast, both moss ADKs were observed as dimers at low concentration (MW of 81 ± 4 kDa) (Figure 2B). Thermostability measurements by nanoDSF revealed similar melting temperatures (*T_m_*) between 53°C and 64°C (Figure S1) for all isoforms, with a minor stabilizing effect of ATP (up to 4°C). Notably, ZmADK2 and PpADK2 exhibited slightly greater stability with a *T_m_* of 64 °C.

The analytical ultracentrifugation (AUC) in sedimentation velocity mode, small-angle X-ray scattering (SAXS), and dynamic light scattering (DLS) measurements revealed differences between the two selected and the most readily available representatives ZmADK2 and PpADK1 (Figures 2C, 2D and 2E). AUC data showed that the dominant component in the ZmADK2 sample is the monomer at such concentration with a sedimentation coefficient value of ∼ 2.8 S. The molecular weight estimate determined using the Svedberg equation 36-39 kDa corresponds to the expected monomer mass of 39 kDa. On the contrary, PpADK1 was monomeric at a low concentration with a sedimentation coefficient of ∼ 2.9 S and dimeric at a higher concentration (over 0.3 mg ml^-1^) with a sedimentation coefficient of ∼ 4.2 S (Figure 2C). Guinier analysis of SAXS data (Figure 2D) using ATSAS software (Manalastas-Cantos et al., 2021) yielded a radius of gyration (R_g_) of 36.77 ± 0.15 Å for PpADK1, in contrast to 22.68 ± 0.02 Å for ZmADK2. Subsequently, the calculated Bayesian MW estimate was 91.2 ± 4.6 kDa and 43.8 ± 3.4 kDa for PpADK1 and ZmADK2, respectively. These estimates are in good agreement with dimeric and monomeric states, respectively.

### Moss ADKs are dimeric upon ATP binding and monomeric upon both ATP and substrate binding

DLS revealed a difference in the hydrodynamic diameter (d_H_) between PpADK1 (d_H_ of 6.9 ± 0.14 nm) and ZmADK2 (d_H_ of 5.0 ± 0.15 nm) (Figure 2E). While the presence of ATP did not affect particle diameter compared with the apoform for both enzymes, the concurrent addition of Ado substrate with ATP induced a shift towards a smaller diameter of PpADK1 (d_H_ of 5.6 ± 0.50 nm). Moreover, incubation with the AP5A inhibitor, mimicking bound Ado with ATP, resulted in an even smaller size of PpADK1 (d_H_ of 5.1 ± 0.12 nm). These data indicate that PpADK1 dimer dissociates upon ternary-complex formation.

### Maize ADKs are more active than moss enzymes

Enzyme activity was first measured using both HPLC (Moffatt et al., 1991) and the coupled reaction assays (Blondin et al., 1994). Later, the coupled reaction was favored because it yielded higher specific activities. We screened the activities with Ado and several cytokinin ribosides at a saturating 100 µM concentration (Table 1). Although all five plant ADKs exhibited the highest activities with Ado, all three maize ADKs were much more active (∼ 30 - 40 nkat mg^-1^) than both moss enzymes (∼ 4-5 nkat mg^-1^). Activities with cytokinin ribosides like iPR, tZR, or 6-benzylaminopurine riboside (BAPR) were comparable within each isoform and up to 10 times lower than those with Ado. Because iPR and *t*ZR are *N*^6^-isopentenylAdo derivatives, we also prepared four other Ado derivatives (Figure S2-S6), shorter than naturally occurring cytokinins. We did the screening with two longer ones, namely *N*^6^-isopropylAdo and *N*^6^-isobutylAdo. They were phosphorylated at rates comparable to those observed for cytokinin ribosides, and no significant variation was observed among the studied ADK isoforms (Table 1). The *K*_m_ values for Ado were in the high nanomolar range (0.5-0.7 µM) for all five ADKs, whereas those for iPR were in the low micromolar range (4-12 µM) (Table 2). Maize ADKs displayed catalytic efficiency (*k*_cat_/*K*_m_) values 50 to 150-fold higher for Ado than for iPR (Table 2), similar to moss ADKs.

**Table 1.** Specific activities of five plant ADKs with several putative nucleoside substrates at 100 µM concentration. Measured in a reaction mixture 50 mM Tris-HCl buffer pH 7.5 containing 50 mM KCl, 6 mM MgCl_2_, 300 µM PEP, 0.5 mM DTT, 200 µM NADH, 10 mM ATP, 2 units of PK and 2.5 units of LDH at 30 °C.

| Enzyme | Specific activity (nmol s<sup>-1</sup> mg<sup>-1</sup>) |  |  |  |  |  |
| --- | --- | --- | --- | --- | --- | --- |
|  | *Ado* | *iPR* | *tZR* | *BAPR* | *N<sup>6</sup>-IsopropylAdo* | *N<sup>6</sup>-IsobutylAdo* |
| ZmADK1 | 30.5 ± 0.4 | 2.8 ± 0.3 | 2.7 ± 0.3 | 3.3 ± 0.2 | 2.2 ± 0.2 | 2.1 ± 0.4 |
| ZmADK2 | 40.3 ± 1.4 | 4.1 ± 0.4 | 3.8 ± 0.4 | 5.1 ± 0.6 | 3.9 ± 0.5 | 4.2 ± 0.6 |
| ZmADK3 | 30.0 ± 1.1 | 5.1 ± 0.3 | 2.7 ± 0.4 | 5.3 ± 0.4 | 3.8 ± 0.3 | 4.8 ± 0.5 |
| PpADK1 | 5.09 ± 0.26 | 0.93 ± 0.11 | 0.74 ± 0.06 | 0.84 ± 0.04 | 0.96 ± 0.09 | 0.90 ± 0.06 |
| PpADK2 | 4.70 ± 0.05 | 0.83 ± 0.08 | 0.83 ± 0.08 | 0.89 ± 0.07 | 0.83 ± 0.05 | 0.75 ± 0.08 |

**Table 2.** Kinetic parameters of plant ADKs from moss and maize for Ado and iPR. Saturation curves were measured by a coupled reaction with PK and LDH in 50 mM Tris-HCl buffer pH 7.5 containing 50 mM KCl, 6 mM MgCl_2_, 300 µM PEP, 0.5 mM DTT, 200 µM NADH, 10 mM ATP at 30 °C. Kinetic constants *K*_m_ and *k*_cat_ were determined using GraphPad Prism 8.0 software (http://www.graphpad.com). Standard deviations were calculated from three repetitive measurements.

| Enzyme | Ado |  |  | iPR |  |  |
| --- | --- | --- | --- | --- | --- | --- |
| | $K_m$<br>$\mu M$ | $k_{cat}$<br>$s^{-1}$ | $k_{cat}/K_m$<br>$M^{-1} s^{-1}$ | $K_m$<br>$\mu M$ | $k_{cat}$<br>$s^{-1}$ | $k_{cat}/K_m$<br>$M^{-1} s^{-1}$ |
| ZmADK1 | $0.62 \pm 0.11$ | $1.14 \pm 0.04$ | $1.8 \pm 0.3 \times 10^3$ | $8.5 \pm 0.7$ | $0.10 \pm 0.002$ | $1.2 \pm 0.4 \times 10^1$ |
| ZmADK2 | $0.65 \pm 0.09$ | $1.55 \pm 0.04$ | $2.4 \pm 0.3 \times 10^3$ | $9.8 \pm 1.5$ | $0.19 \pm 0.007$ | $2.0 \pm 0.3 \times 10^1$ |
| ZmADK3 | $0.51 \pm 0.04$ | $1.18 \pm 0.03$ | $2.3 \pm 0.8 \times 10^3$ | $4.3 \pm 0.4$ | $0.22 \pm 0.007$ | $5.0 \pm 1.8 \times 10^1$ |
| PpADK1 | $0.71 \pm 0.05$ | $0.21 \pm 0.01$ | $3.0 \pm 0.7 \times 10^2$ | $12.1 \pm 2.5$ | $0.03 \pm 0.002$ | $2.5 \pm 0.6$ |
| PpADK2 | $0.54 \pm 0.05$ | $0.18 \pm 0.01$ | $3.3 \pm 0.8 \times 10^2$ | $4.0 \pm 0.3$ | $0.03 \pm 0.001$ | $7.3 \pm 0.4$ |

Binding affinities (*K*_D_ values) for the tested substrates such as cytokinin ribosides (transport form of the hormone) were further determined by microscale thermophoresis (MST) with the two selected representatives, ZmADK2 and PpADK1 (Figure 2F and 2G, Table 3), displaying a similar binding pattern with *K*_D_ values close to *K*_m_ values. The best affinity was observed for Ado (*K*_D_ of 0.4-0.7 µM), followed by iPR, *t*ZR and BAPR with *K*_D_ values ranging between 5 to 25 µM. Interestingly, the large molecule of AP5A inhibitor was bound in two subsequent steps, the first *K*_D_ value corresponding to its binding into the Ado site and the second one corresponding to its binding into the ATP site. For PpADK1, the lower *K*_D_ value was 8.1 ± 1.1 µM, while the higher *K*_D_ value was 0.63 ± 0.12 mM. Affinities for the cosubstrate ATP and its product ADP were relatively weak and in the low millimolar range. The measured *K*_D_ values for ATP were 2.2 ± 0.6 mM with ZmADK2 and 2.9 ± 0.7 mM with PpADK1, respectively.

**Table 3.** Binding affinities and *K*_m_ values of several Ado and ATP analogs with ZmADK2. Activities were measured in a coupled reaction with PK and LDH at 30 °C in 50 mM Tris-HCl buffer pH 7.5 using 10 mM ATP. Affinity was measured by MST in 50 mM HEPES buffer pH 7.5, 1 mM MgCl2, and 0.1% Tween. n.d. - not determined, * - second *K*_D_ value.

| Nucleoside/nucleotide ligand | $K_D$<br>$\mu M$ | $K_m$<br>$\mu M$ | $k_{cat}$<br>$s^{-1}$ |
| --- | --- | --- | --- |
| Ado | 0.40 ± 0.14<br>1480 ± 230* | 0.65 ± 0.09 | 1.55 ± 0.04 |
| *N*<sup>6</sup>-MethylAdo | 8.7 ± 2.5 | 10.7 ± 2.2 | 0.24 ± 0.02 |
| *N*<sup>6</sup>,*N*<sup>6</sup>-DimethylAdo | 12.3 ± 4.6 | 11.1 ± 2.5 | 0.20 ± 0.02 |
| *N*<sup>6</sup>-IsopropylAdo | 52.8 ± 13.9 | 54.8 ± 6.8 | 0.19 ± 0.01 |
| *N*<sup>6</sup>-IsobutylAdo | 45.9 ± 9.3 | 70.4 ± 8.8 | 0.21 ± 0.01 |
| iPR | 4.8 ± 1.1 | 9.8 ± 1.5 | 0.19 ± 0.01 |
| 2'-DeoxyAdo | 505 ± 94 | 502 ± 84 | 0.45 ± 0.02 |
| 3'-DeoxyAdo (cordycepin) | 175 ± 9 | 178 ± 62 | 0.44 ± 0.02 |
| 7-DeazaAdo (tubercidin) | 2.2 ± 0.7 | 2.5 ± 1.5 | 0.17 ± 0.01 |
| AraA (vidarabine) | >> 500, n.d. | 560 ± 115 | 0.24 ± 0.02 |
| Inosine | 440 ± 180 | 395 ± 50 | 0.57 ± 0.03 |
| Guanosine | 217 ± 107 | 184 ± 61 | 0.46 ± 0.04 |
| Xanthosine | 166 ± 77 | 187 ± 31 | 0.55 ± 0.02 |
| Uridine | >> 5000, n.d. | >> 1000, n.d. | 0.15 ± 0.04 |
| Cytidine | 286 ± 10 | 307 ± 51 | 0.23 ± 0.01 |
| AP5A | 14.5 ± 9.2<br>949 ± 46* | - | - |
| AMP | 247 ± 71<br>8299 ± 2070* | n.d. | n.d. |
| ADP | 10300 ± 1190 | n.d. | n.d. |
| ATP | 2200 ± 620 | n.d. | n.d. |

We analyzed the binding properties of several other purine ribosides, deaza, and deoxy derivatives with the most active enzyme, ZmADK2 (Table 3, Figure S1). None of the ligands reached *k*_cat_ values or had as high affinity as Ado. The four Ado-derivatives, namely *N*^6^-methylAdo, *N*^6^, *N*^6^-dimethylAdo, isopropylAdo and *N*^6^-isobutylAdo, were synthesized to analyze the effect of side chain on the activity and binding. While *k*_cat_ values were comparable, *N*^6^-methylAdo and *N*^6^, *N*^6^-dimethylAdo showed a slightly lower affinity than iPR, whereas the two longer compounds displayed a 10-fold weaker affinity. iPR. Similarly to AP5A, Ado with AMP displayed two binding events, suggesting they can bind into both the substrate and ATP sites. Ado and AMP exhibited the second binding event in the low millimolar range (*K*_D(2)_ values of 1.48 ± 0.23 mM and 8.3 ± 2.0 mM, respectively) for their binding in the ATP site.

AraA proved to be a very poor ligand, with a *K*_D_ value exceeding 500 µM, making it challenging to measure due to solubility issues. Both 2′-deoxy Ado and 3′-deoxy Ado derivatives displayed roughly 1200-fold and 400-fold weaker affinity compared to Ado, with *K*_D_ values of 505 ± 94 µM and 175 ± 9 µM, respectively (Table 3). This underscores the crucial role of both O2’ and O3’ hydroxyl groups in nucleoside binding. In contrast, binding of the 7-deazaAdo derivative was significantly less affected, displaying *K*_D_ value of 2.2 ± 0.7 µM, which is ∼ 5.5-fold weaker affinity than Ado. ZmADK2 could also bind and phosphorylate oxopurine nucleosides such as inosine, xanthosine or guanosine. Measured *k*_cat_ values were around one-third of what was observed with Ado. *K*_D_ and *K*_m_ values fell within the high micromolar range, indicating that these compounds are relatively weak substrates.

### The crystal structure of maize ADK2 reveals a unique dimer formation among plant ADKs

Among the two selected representatives, ZmADK2 and PpADK1, we only obtained crystals of the apo form of ZmADK2 and solved the structure (PDB 8RF7) by molecular replacement using the structure of the human ADK in the open conformation (PDB 2I6B, Muchmore et al., 2006) as a search model (Table S1). ZmADK2 shares ∼ 58 % sequence identity with human ADK (HsADK, Figure S7).

Crystallization conditions of ZmADK2 favored the dimeric form to be present in the asymmetric unit (Figures 3A and 3B), which contains two ZmADK2 molecules, each in the open conformation, and resembles that of HsADK with an average root mean square (RMSD) of ∼ 1.2 and 1.4 Å for all Cα atoms. Each monomer is composed of two domains: a large ten-stranded α/β Rossmann-like nucleotide-binding domain (residues 1-14, 64-117 and 137-340) comprising the ATP binding site and a small five-stranded α/β domain (residues 17-61 and 120-134). The small domain encompases β2, β3, β4, β8 and β9 sheets plus α1 and α2 helices. Both domains are connected *via* four peptide hinges (Leu15-Leu16, Gly62-Gly63, Thr118-Gly119, Asn135-Leu136). The nucleoside substrate pocket is located in a cleft between the two domains.

**Figure 3.**
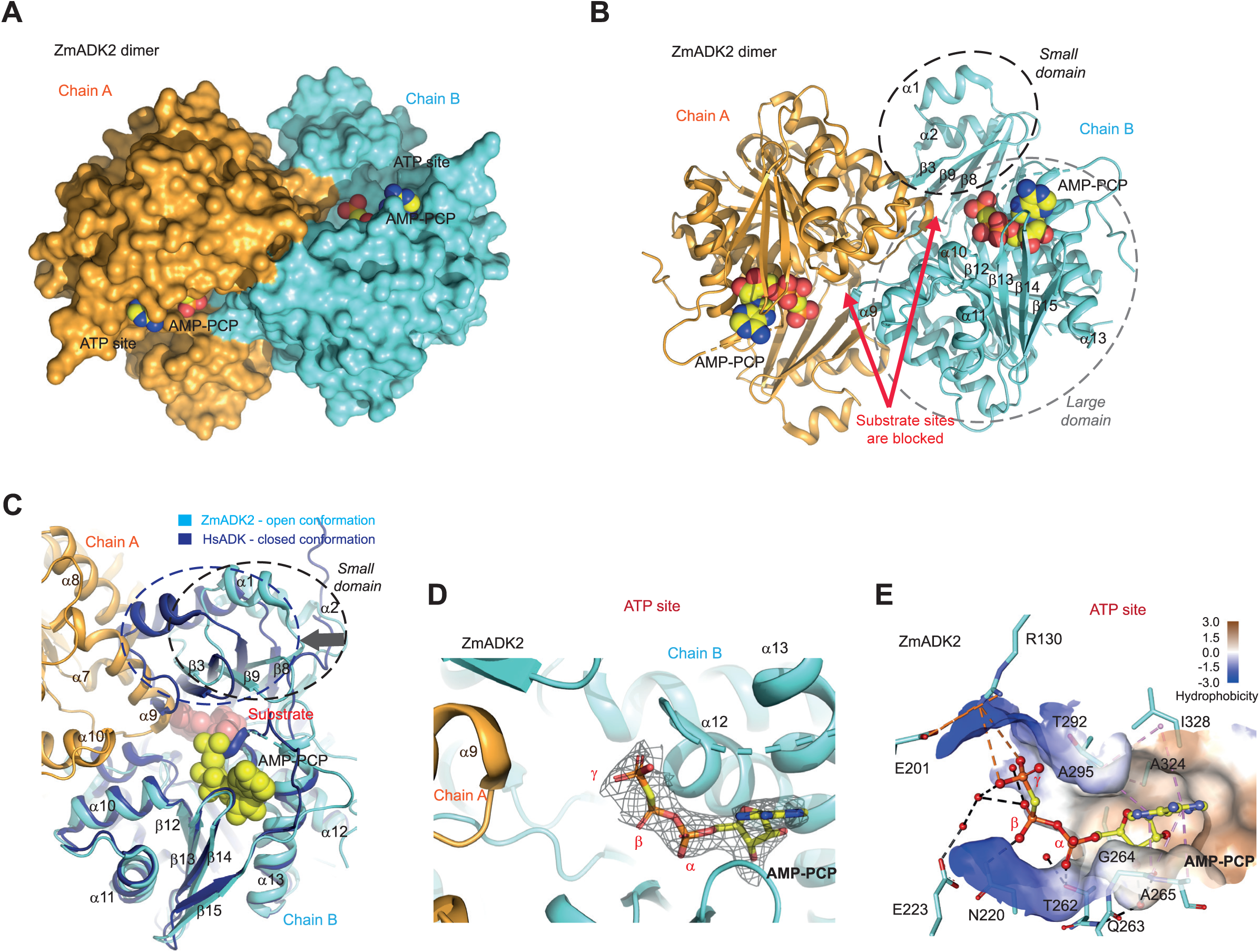
Dimeric structure of plant maize ADK2 in open conformation. **(A)** Dimeric structure of ZmADK2 (this work, PDB 8RGJ) in surface representation colored in cyan/orange with the bound AMP-PCP in yellow. **(B)** Cartoon representation of the dimeric structure of ZmADK2 with bound AMP-PCP, featuring a topology description. Both small and large domains are labeled. Red arrows indicate the position of each substrate-binding site blocked by the neighboring subunit. **(C)** Superposition of ZmADK2 in the open conformation (chain B, cyan) with HsADK in the closed conformation (PDB 26IA, dark blue, Muchmore et al., 2006) containing bound nucleoside. The black arrow denotes the rotation of the small domain. Chain A of the ZmADK2 dimer is in orange. **(D)** Binding of the AMP-PCP in the ATP site of ZmADK2 in its annealing Fo-Fc omit map (black mesh) contoured at 2.5 σ. **(E)** Binding interactions of AMP-PCP in ZmADK2 and hydrophobicity of the ATP site. Hydrogen bonds are shown in black dashed lines, while electrostatic and hydrophobic interactions are shown as orange and pink dashed lines.

The dimer interface covers an area of 1577 Å^2^ as calculated by the PISA server (www.ebi.ac.uk/pdbe/pisa/) involving about 50 residues per monomer and making it characteristic of biological interactions (Janin et al., 2007) (Figure S7). The solvation-free energy gained upon interface formation is −19.7 kcal mol^-1^. This involves side chains of both domains: the regions 35-43 and 130-136 of the small domain and the regions 142-145, 167-174, 196-206 and 222-236 of the large domain. So far, such a dimer arrangement has not been observed for any ADKs.

Substrate binding induces the closed conformation, resulting in a rotation of the small domain by approximately 30°, as documented in previous studies (Mathews et al., 1998; Muchmore et al., 2006). Remarkably, the dimer interface prevents displacement of the small domains. Indeed, the superposition of HsADK in the closed conformation (PDB 1BX4, Mathews et al., 1998) on one subunit of ZmADK2 dimer showed that helices α8 and α9 of the large domain of the other subunit block the rotation of its small domain (α1-β3-α2 segment) and thus proper binding of the nucleoside substrate (Figure 3C). Therefore, to be active, moss and maize ADK dimers must be dissociated into monomers.

### ATP binding has no effect on the dimeric state

In agreement with DLS measures showing that ATP binding did not affect the oligomeric state of plant ADKs, soaking crystals of apo ZmADK2 dimer with the non-hydrolyzable ATP analog lead to a dimeric complex with AMP-PCP (PDB 8RGJ) occupying the ATP binding site. Its γ-phosphate moiety is oriented toward the substrate binding site where the ribose moiety of a nucleoside substrate would be located (Figures 3D and 3E). The Ado moiety of ATP is surrounded by hydrophobic residues. Indeed, its ring is located between Ile328 and Ala295 on one side, and Ala265 with Gly264 on the other, forming multiple hydrophobic contacts (Figure 3E). The O2’ atom of ribose moiety is bound directly to the SG atom of Cys321, and the O3’ atom is H-bonded *via* a water molecule to the main-chain oxygen atom of Gly264 and nitrogen of Gln263. The α-phosphate group forms a hydrogen bond with the OG1 atom of Thr262. The β-phosphate group is H-bonded to Asn220, and both the β- and γ-phosphate groups interact with Glu223 *via* two water molecules. Finally, the negatively charged γ-phosphate moiety establishes an electrostatic interaction with the positively charged side-chain of Arg130.

The role of Asn220 and Glu223 residues was studied by site-directed mutagenesis on ZmADK3 (Table 4). The N222A variant (ZmADK3 numbering is shifted by two amino acids) showed a 7-fold lower affinity for ATP with half activity compared to wild-type (WT). Remarkably, the E225A variant displayed approximately a 6-fold lower affinity for ATP and only 6% activity compared to WT.

**Table 4.**
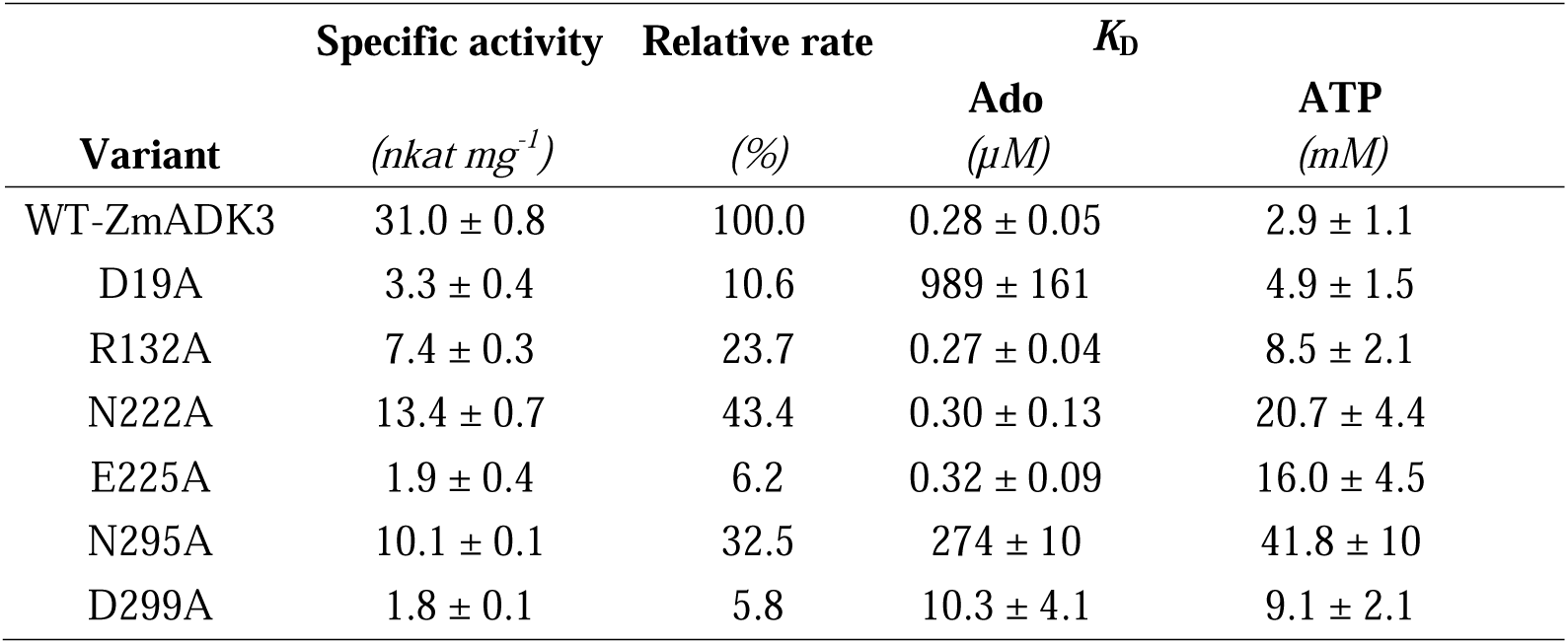
Characterization of active-site variants of ZmADK3. Activities were measured with 100 µM Ado in a coupled reaction with PK and LDH at 30 °C in 50 mM Tris-HCl buffer pH 7.5 using 10 mM ATP. *K*_D_ values were measured by MST in 50 mM HEPES buffer pH 7.5, 1 mM MgCl_2_ and 0.1% Tween.

| Variant | Specific activity | Relative rate | $K_D$ | |
| --- | --- | --- | --- | --- |
| | (*nkat mg<sup>-1</sup>*) | (%) | Ado<br>(μM) | ATP<br>(mM) |
| WT-ZmADK3 | 31.0 ± 0.8 | 100.0 | 0.28 ± 0.05 | 2.9 ± 1.1 |
| D19A | 3.3 ± 0.4 | 10.6 | 989 ± 161 | 4.9 ± 1.5 |
| R132A | 7.4 ± 0.3 | 23.7 | 0.27 ± 0.04 | 8.5 ± 2.1 |
| N222A | 13.4 ± 0.7 | 43.4 | 0.30 ± 0.13 | 20.7 ± 4.4 |
| E225A | 1.9 ± 0.4 | 6.2 | 0.32 ± 0.09 | 16.0 ± 4.5 |
| N295A | 10.1 ± 0.1 | 32.5 | 274 ± 10 | 41.8 ± 10 |
| D299A | 1.8 ± 0.1 | 5.8 | 10.3 ± 4.1 | 9.1 ± 2.1 |

### Closed conformation is associated with the monomeric state and ternary complex formation

Co-crystallization of ZmADK3 and PpADK1 in the presence of AP5A and a mixture of Ado and ADP, respectively, led to two high-resolution crystal structures of a ternary complex (PDB 8RPA and 9FW6; Table S1, Figures 4A and 4B). The asymmetric unit contains one monomer for the ZmADK3-AP5A complex (Figure 4A) and two monomers for the PpADK1-Ado-ADP complex, all in the closed conformation. Structural comparison between ZmADK2 (open state) and the closed state of both ZmADK3 and PpADK1 reveals that AP5A (or Ado) binding induced a rigid-body motion of the smaller domain towards the large domain, resulting in the closed conformation. Two helices, α1 and α2, shift by up to 12 Å to shield the Ado site from the solvent (Figures 4C and 4D).

**Figure 4.**
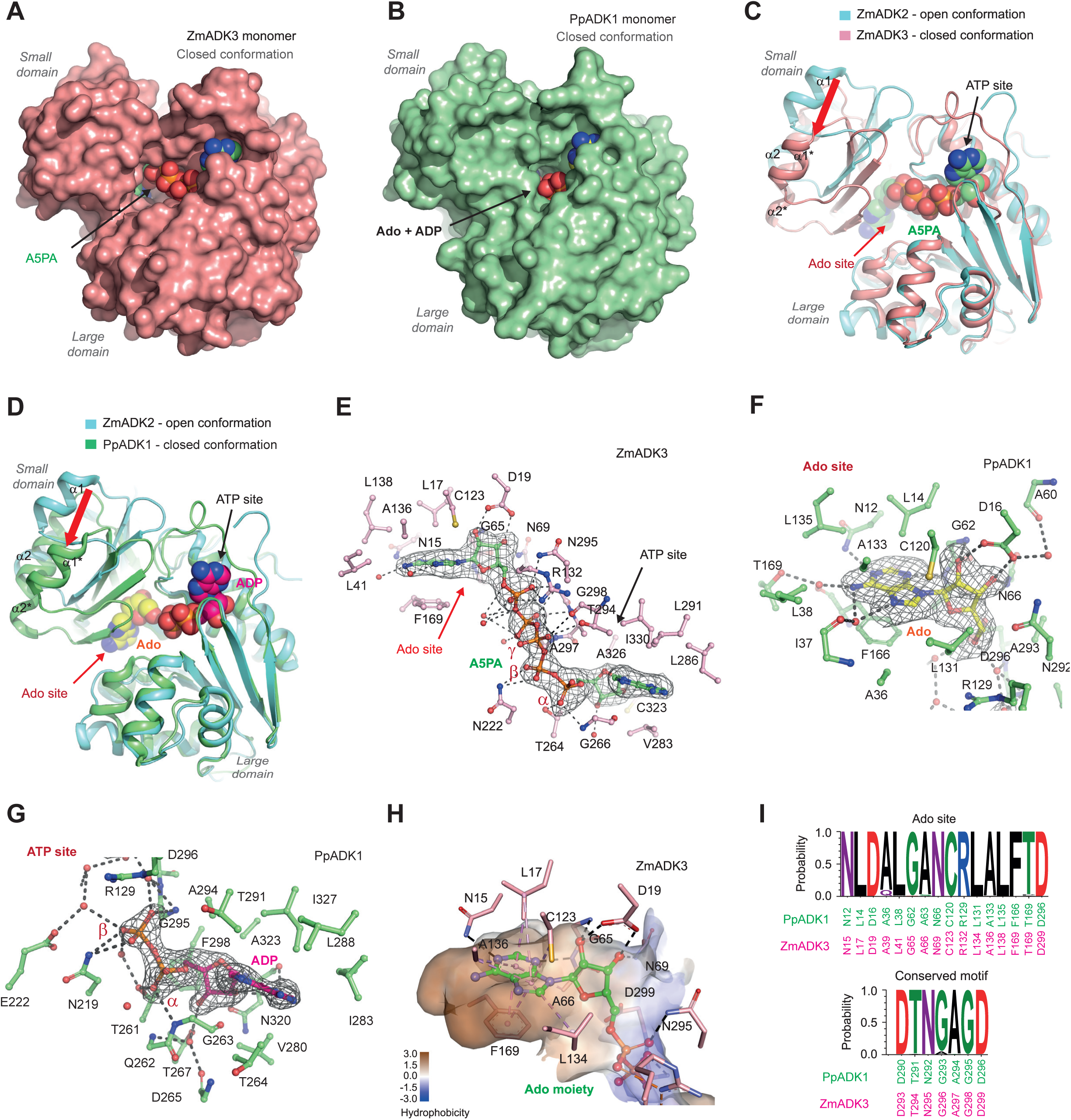
Monomeric structure of plant ADK in closed conformation. **(A)** The monomeric structure of ZmADK3 (this work, PDB 8RPA) colored in pink with the bound AP5A in green. **(B)** The monomeric structure of PpADK1 (this work, PDB 9FW6) colored in green with both bound Ado and ADP. **(C)** Superposition of ZmADK2 in an open conformation (cyan) with ZmADK3 with AP5A in the closed conformation (pink). The red arrow indicates the shift of the small domain. **(D)** Superposition of ZmADK2 in an open conformation (cyan) with PpADK1 with Ado and ADP in the closed conformation (green). The red arrow indicates the shift of the small domain. **(E)** Interactions between ZmADK3 and the bound AP5A inhibitor. AP5A (in green) is shown in its annealing Fo-Fc omit map (black mesh) contoured at 3.0 σ. **(F)** Interactions of Ado ligand, colored in yellow, in the Ado binding site of PpADK1 with the enzyme. The Ado is shown in its annealing Fo-Fc omit map (black mesh) contoured at 3.0 σ. **(G)** Binding of the ADP ligand in the ATP site of PpADK1 in its annealing Fo-Fc omit map (black mesh) contoured at 3.0 σ and its interactions with the enzyme. **(H)** Hydrophobic and hydrogen-binding interactions in the adenosine pocket of ZmADK3 with AP5A ligand. **(I)** The conservation of amino acid residues in plant ADKs forming the Ado site and conserved motif involved in binding of the β- and γ-phosphate of ATP. The sequence logo was made using WebLogo 3 (http://weblogo.threeplusone.com).

The electron density map of the bisubstrate analog AP5A in ZmADK3 is very well defined (Figure 4E). The ligand binds in the groove between the large and small domains. The electron density maps of Ado and ADP in PpADK1 show clearly that both substrates bind in the Ado and ATP binding sites, respectively (Figures 4F and 4G). When superposed, Ado in PpADK1 overlaps the Ado part of AP5A in a highly hydrophobic pocket (Figure 4H). Analysis of numerous plant sequences revealed a high conservation of residues forming the Ado pocket, as illustrated in Figure 4I. This finding explains the remarkably similar substrate properties observed among the studied ADK isoforms. The unique DTN(G/A)AGD motif near the C-terminus (Figure S7), typical for ADKs, is fully conserved and comprises residues involved in the binding of β- and γ-phosphates of ATP.

The purine ring of Ado stacks on the phenyl ring of Phe169 (in ZmADK3 numbering; PpADK1 numbering is shifted by three amino acids) and establishes hydrophobic interactions with Leu17, Ala66, Cys123, Leu134 and Ala136. The N1 and N3 atoms of the purine ring are H-bonded to the side chain of Asn15 and the main-chain nitrogen atom of Ala66, respectively. Both O2’ and O3’ atoms of the ribose moiety bind to the side chain of Asp19. In addition, the O3’ atom interacts with Asn69, while the O2’ atom is H-bound to the main-chain nitrogen atom of Gly65. The phosphate group, transferred to Ado to form AMP, interacts with the side chains of Asn295 and Arg302.

AP5A in ZmADK3 and AMP-PCP in ZmADK2 adopt the same conformation and position in the ATP binding site. They establish similar hydrogen and hydrophobic interactions. The adenine part of ADP in PpADK1 is slightly shifted compared with the adenine moiety of the ATP part of AP5A due to the presence of a threonine at position 264 in PpADK1 instead of an alanine (Ala265) in ZmADK3. The γ-phosphate in AP5A forms a hydrogen bond with the OG1 atom of Thr294 and the side chain of Arg132.

The crucial role of Asp19 in riboside binding was further confirmed by site-directed mutagenesis (Table 4). The D19A variant exhibited a 10-fold reduction in specific activity and a poor affinity for Ado. The *K*_D_ value for Ado shifted from 0.28 µM to 989 µM concentration. Although Asp299 is placed behind the ribose of Ado moiety of AP5A with no direct H-bonds (Figure 3I), it would bind the free O5’ hydroxyl group. The D299A variant displayed only ∼ 6% activity and approximately 37 times lower affinity for Ado than WT. Also, the Arg residue (Arg130 in ZmADK2 and Arg132 in ZmADK3), situated between Ado and ATP pockets, plays a vital role in catalysis. This residue holds the γ-phosphate of ATP, as seen in the ZmADK2 structure (Figure 3E), and the phosphate molecule bound to Ado during AMP formation, as observed in the ZmADK3 structure (Figure 4E). The R132A variant displayed ∼ 24% activity compared to the WT, and while the affinity to Ado remained unaffected, the affinity to ATP was three times lower.

Finally, *in silico* docking of cytokinin riboside (iPR) into the Ado site of ZmADK3/PpADK1 revealed enough space to accommodate the isoprenoid chain of cytokinin next to the side chains of L41/L38 and F200/F197, which can adopt different rotamer orientations. The position of iPR overlaps that of Ado (Figure S8), and the binding energies of both ligands were nearly identical (Table S3).

### All ADK genes are highly expressed in maize, but only one in moss

The expression of the three maize ADK genes was observed in roots and leaves during early developmental stages, as assessed by qPCR using FAM-TAMRA probes (Figure 5A, Table S4). In young seedlings, *ZmADK2* transcripts were highly abundant in stems, leaves and roots, while *ZmADK3* transcripts were primarily found in leaves and *ADK1* transcripts were observed in roots. In older plants, *ZmADK2* expression remained high across various samples, including leaves, roots, tassels, silks, and kernels. Transcripts of *ZmADK3* were abundant in silks and kernels, while *ZmADK1* transcripts were mainly present in kernels. This expression pattern suggests a vital role of all three ADKs in developing seeds and reproductive organs containing dividing cells. Under salt stress, the expression of *ZmADK1* and *ZmADK2* increased in roots. Treatment with active cytokinin (*t*Z) did not induce significant changes in expression levels in either leaves or roots. Although nitrogen starvation is known to upregulate *ENT* genes and lead to increased nucleoside import from degraded RNAs (Cornelius et al., 2012; Melino et al., 2018), the expression of *ADK* genes was not upregulated upon nitrogen starvation in maize.

**Figure 5.**
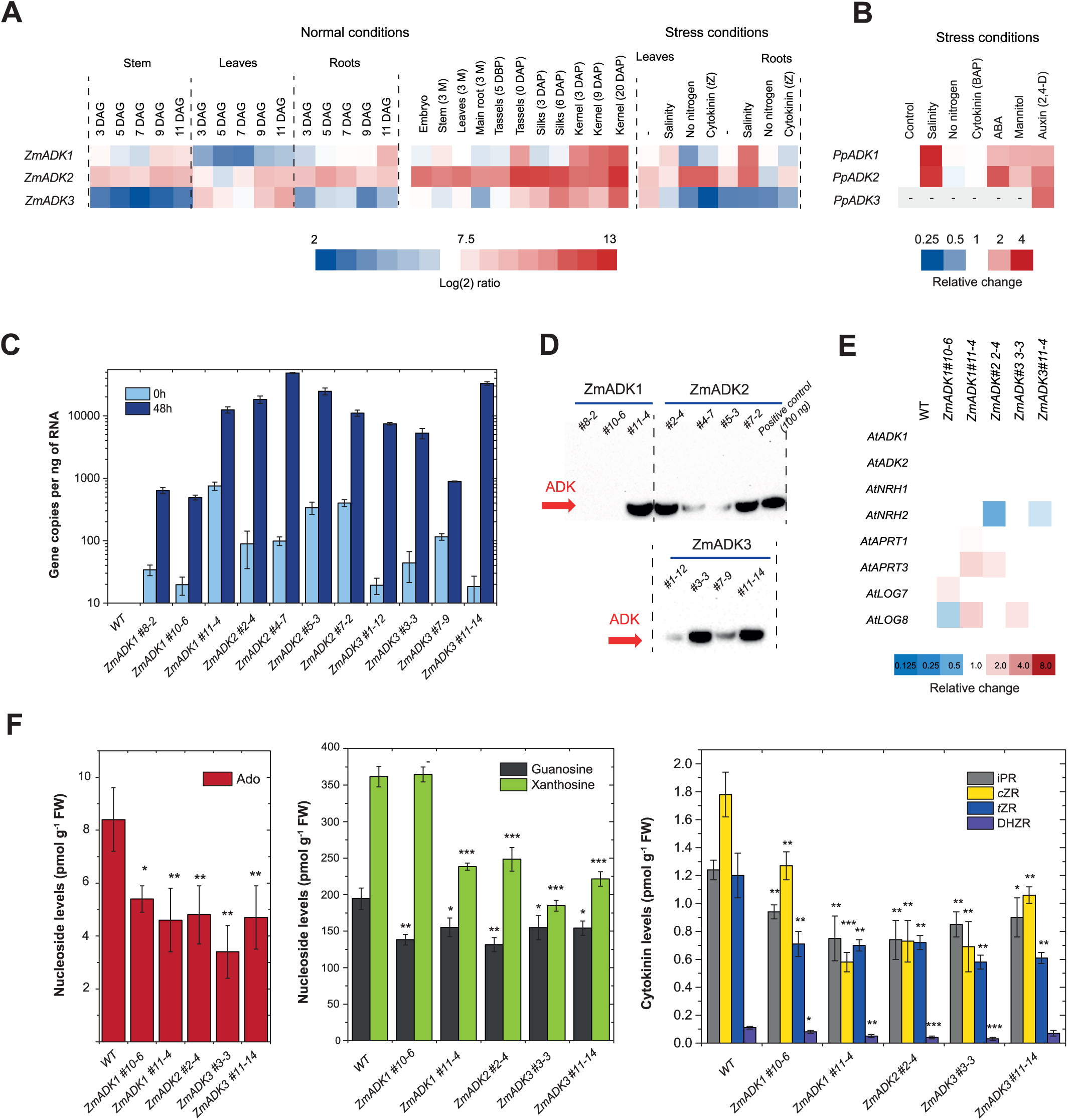
Expression of *ADKs* in maize and moss and characterization of *ZmADK* transgenic lines. **(A)** *ADK* expression in maize was followed for two weeks after germination and later in various organs under normal conditions (left side) and during various stresses (right side). The heat map illustrates transcript levels detected in 1 ng of total RNA. All values are expressed as log_2_-ratios. DAG, days after germination; DAP, days after pollination; DBP, days before pollination; M, months. **(B)** *ADK* expression in moss grown in liquid Knop’s medium and then exposed to 10 μM abscisic acid, cytokinin (BA), auxin (2,4-D) or methyl jasmonate, 200 mM NaCl or 400 mM mannitol for 4 days. **(C, D)** Dexamethasone-inducible expression of *ZmADK* genes in homozygous Arabidopsis seedlings after 48 hours at 14 DAG, assessed at RNA **(C)** and protein levels **(D)**. The expression was tested using qPCR with FAM-TAMRA probes and protein production was evaluated from whole plantlet extracts via Western blot. The PVDF membrane was probed with a monoclonal anti-6His-tag mouse antibody and reacted with mouse IgG BP-HRP conjugate for chemiluminescent detection. **(E)** The expression of related genes as analyzed by qPCR using labeled probes upon 48-hour dexamethasone induction of *pOpON::ZmADK* transgenic lines at 14 DAG. **(F)** Nucleoside levels (pmol g^-1^ FW) in induced *pOpON::ZmADK* overexpressor lines including Ado, guanosine, xanthosine and cytokinin ribosides. Measured in three technical replicates. Asterisks indicate statistically significant differences in induced transgenic lines versus WT-Col0 in a Student’s t-test (t-test; *, **, and *** correspond to P-values of 0.05 > p > 0.01, 0.01 > p > 0.001, and p < 0.001, respectively).

In *Physcomitrella patens* (protonemal stage) cultivated in liquid Knop medium, we observed that *PpADK1* transcripts were highly abundant, reaching approximately 13800 copies ng^-1^ of RNA (Table S5). Conversely, the amount of *PpADK2* transcripts was two orders of magnitude lower (∼ 80 copies ng^-1^ of RNA) and transcripts of *PpADK3* were not detectable. When exposed to salt stress, the expression of both *PpADK1* and *PpADK2* was roughly 3-fold higher than in the control (Figure 5B). Additionally, abscisic acid and mannitol treatments increased the expression of both genes approximately twofold. On the contrary, treatment with active cytokinin (BAP) or nitrogen starvation did not induce significant changes in *ADK* expression levels, mirroring our findings in maize. The only condition in which the *PpADK3* transcripts were detected occurred after moss treatment with the synthetic auxin 2,4-dichlorophenoxyacetic acid (2,4-D).

In summary, the genome of *Physcomitrella* lacks the *PNP* gene; *PpADK2* and *PpADK3* genes are poorly expressed, and both tested moss enzymes, PpADK1 and PpADK2 display weak activities compared to maize ADKs. All these findings imply that the pathway from Ade to AMP via Ado is rather suppressed, and the direct conversion from Ade to AMP via APTs plays a significant role in *Physcomitrella*.

### ZmADK overexpressors contain lower levels of Ado and cytokinin ribosides

*Arabidopsis thaliana* Col0 plants were stably transformed with expression constructs containing cDNAs of three maize ADKs placed under the control of a dexamethasone-inducible promoter. Independent T3 homozygous transgenic lines were selected for each overexpression construct according to the induced gene expression level identified by qPCR (Figure 5C). Transcript levels of *ZmADKs*, measured 72 h after induction, displayed significant variations among the studied lines. At least three lines, namely *pOpOn::ZmADK1*#10-6, *ZmADK1*#8-2 and *ZmADK3*#7-9, had a lower amount of transcript copies (below 1000 per ng of RNA) than the others (above 10000 copies). Similarly, the above lines accumulated a lower amount of ZmADKs at the protein level. On the other hand, five other lines contained much higher ZmADK1/2/3 levels (Figure 5D). We selected both representatives with weak and strong *ZmADK* overexpression for further characterization *in planta*.

The qPCR analysis conducted on all examined transgenics (Figure 5E) revealed no significant changes two days after induction in the expression of the two Arabidopsis *ADK* genes, namely *AtADK1* (At3g09820) and *AtADK2* (At5g03300), as well as of both *NRH* genes, namely *AtNRH1* (At2g36310) and *AtNRH2* (At1g05620). Additionally, a subtle upregulation of *AtAPT3* (At4g22570) was observed in two transgenic lines, while no significant changes were detected for *AtAPT1* (At1g27450). Two *pOpOn::ZmADK* lines showed a minor upregulation of the *AtLOG8* (At5g11950) and one line an upregulation of *AtLOG7* (At5g06300) expression. Taken together, no clear pattern in the expression of these genes was observed.

Quantitative analysis of purine and cytokinin bases, nucleosides, and nucleotides in 2-week-old and dexamethasone-induced seedlings was conducted using UPLC-MS/MS as outlined in prior studies (Kopečná et al., 2013). In all analyzed transgenic lines, there was a discernible impact on purine and cytokinin metabolism, as shown in Figure 5F and detailed in Table S6. These alterations were consistent with the substrate preferences of ZmADKs, leading to decreased levels of various nucleosides. Specifically, levels of Ado were reduced to approximately 45 - 65%, while cytokinin ribosides iPR, tZR, cZR, and DHZR were decreased to around 50-70% compared to WT-Col0 plants. Adenine levels were slightly reduced and levels of cytokinin bases remained unchanged. Although AMP levels could not be adequately assessed due to the lack of an internal standard, the levels of cytokinin monophosphates iPRMP, tZRMP, cZRMP, and DHZRMP were not elevated; instead, they were similar or slightly lower (by about 10%) compared to WT. In addition to these changes, among other purine nucleosides, guanosine levels were also slightly reduced to approximately 80%, and xanthosine levels decreased to about 50-65% in the *pOpOn::ZmADK* transgenics compared to WT. Among other cytokinin metabolites, no changes were observed among *N*^7^-glucosides, *N*^9^-glucosides or *O*-glucosides (not shown).

### Larger roots are observed in ZmADK overexpressors under nitrogen-limiting conditions

Understanding that nitrogen starvation induces RNA catabolism (Kraft et al., 2008), we conducted phenotyping experiments on the Arabidopsis lines overexpressing *ZmADK1/2/3*, explicitly focusing on root and rosette growth. The induced *pOpON::ZmADK* lines exhibited noticeable variations in root growth under nitrogen-starving conditions, including slightly longer primary roots and an increased number of lateral roots (Figure 6A). Thus, we performed a detailed screening involving four transgenic *ZmADK* lines (using 24 biological replicates per line) spanning a two-week period to obtain relevant data. Plants were cultivated under standard nitrogen conditions and nitrogen starvation. Detailed statistics for total root and leaf area are given in Table S7 and Table S8.

**Figure 6.**
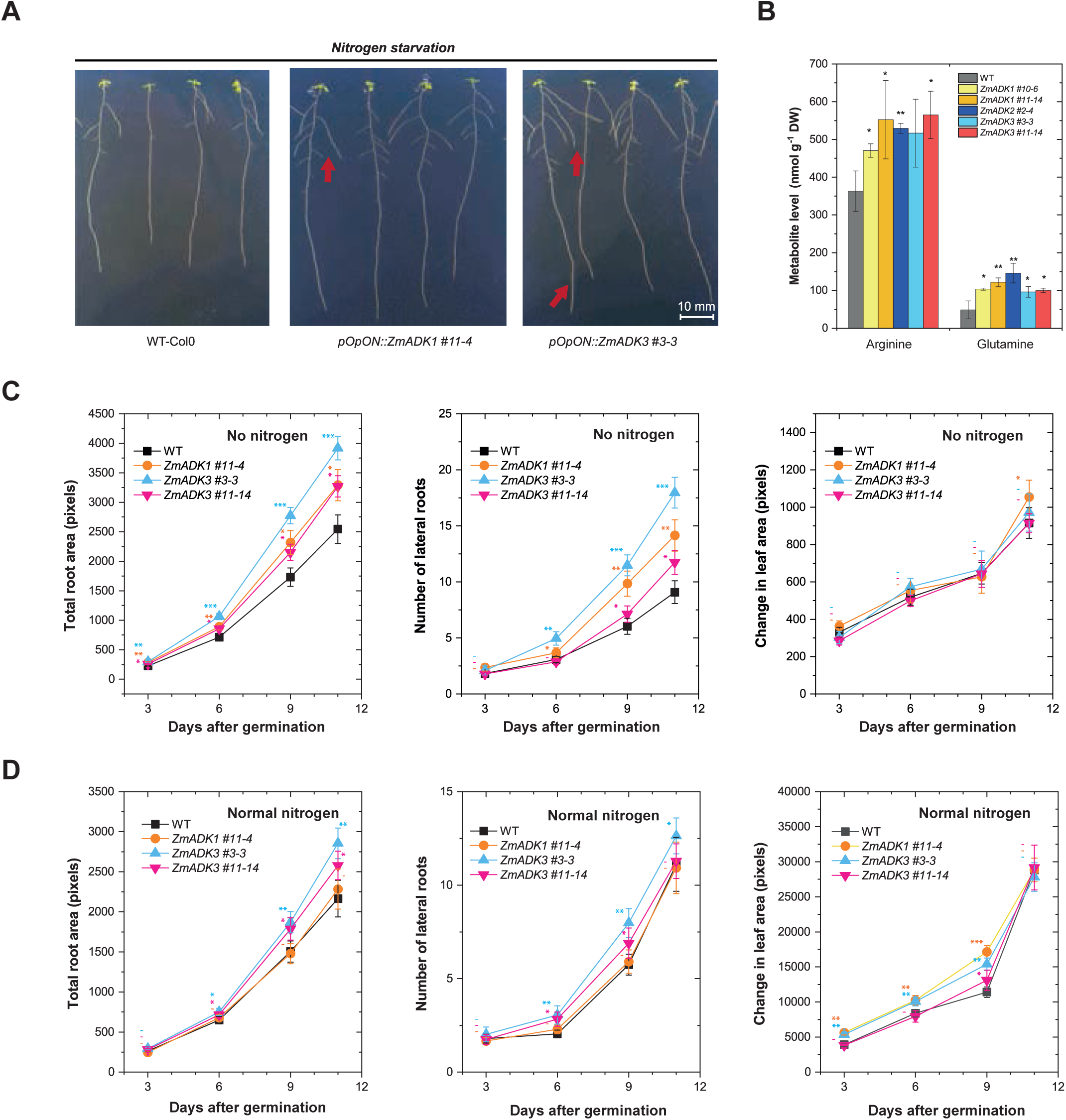
Phenotyping of *A. thaliana pOpON::ZmADK* overexpressors under abiotic stress. **(A)** An illustration of primary and lateral root growth in WT and *pOpOn::ZmADK3* transgenics over 9 days on MS medium without nitrogen. **(B)** Levels of arginine and glutamine in induced *pOpON::ZmADK* overexpressor lines upon nitrogen starvation. Asterisks indicate statistically significant differences in a Student’s t-test (t-test; *, **, and *** correspond to P-values of 0.05 > p > 0.01, 0.01 > p > 0.001, and p < 0.001, respectively). **(C)** Growth of nitrogen-starved plants showing the number of lateral roots, total root area and total rosette area. **(D)** Growth of plants cultivated on standard MS media for up to 14 days. Each experiment included a minimum of 24 seedlings (biological replicates) per *ZmADK* line and growth condition. Asterisks indicate statistically significant differences in transgenic lines compared to WT controls, determined in a paired Student’s t-test (t-test; *, **, and *** correspond to P-values of 0.05 > p > 0.01, 0.01 > p > 0.001, and p < 0.001, respectively).

Notably, nitrogen-starved transgenic plants exhibited a statistically significant increase in the number of lateral roots and total root area (quantified in pixels) (Figure 6C). At 11 days after germination (DAG), mean values for the number of lateral roots in all *ZmADK* transgenics were 30% to 90% higher than those of the wild type (WT). Similarly, mean values for the total root area in ZmADK1#11-4, ZmADK3#3-3, and ZmADK3#11-14 transgenic lines were from 30% to 50% higher than in the WT, with ZmADK2 #2-4 behaving similarly to the WT. In contrast, only minor differences in rosette size were observed, ranging from 89% to 116% compared to WT plants (Figure 6C). Under normal nitrogen conditions, the differences between the induced *ZmADK* transgenics and WT were less apparent (Figure 6D).

Finally, to assess the nitrogen released from purines under nitrogen starvation, we analyzed levels of six free amino acids including ornithine, Arg, Glu, Gln, Asp asn Asn (Table S9). The induced overexpressor lines contained a higher amount of two nitrogen-rich amino acids Arg and Gln (both by ∼ 50%, Figure 6B). While Gln joins the purine catabolism with the Krebs cycle via Glu, Arg stands at the beginning of polyamine synthesis.

## DISCUSSION

### Plant ADKs exist as monomers and dimers

This work presents a comprehensive study of plant ADKs from maize and moss combining *in vivo* and *in vitro* approaches. We showed that plant ADKs can form dimers unlike all other homologous ADKs studied so far (Guranowski, 1979; Andres and Fox, 1979; Miller et al., 1979b; Darling et al., 1999; Mathews et al., 1998; Muchmore et al., 2006; Kuettel et al., 2011; Timm et al., 2014; Schumacher et al., 2000; Zhang et al., 2007; Romanello et al., 2013). *M. tuberculosis* ADK (MtADK) folds into a dimer not comparable to the ZmADK2 dimer, in agreement with a low 22% sequence identity between both enzymes (Figure S9, Long et al., 2003; Reddy et al., 2007). Its structure contains two additional helices in the small domain, which participate in the interface to form the active dimer (Reddy et al., 2007).

We further showed that the ADK dimer can appear only when both subunits are in open conformation as observed in ZmADK2 structure. It cannot be catalytically active as the rotation of the small domain and closure of the substrate cavity for catalysis is impossible. To form the ternary complex and bring close together the two substrates, the enzyme must be monomeric to adopt the closed conformation as shown by the structures of ZmADK3 bound to A5PA and PpADK1 bound to ADP and Ado. DLS on PpADK1 alone or in the presence of ATP with Ado or AP5A highlighted changes in the hydrodynamic diameter of the enzyme upon ligand addition. Indeed, the diameter of PpADK1 (dimeric in solution) became smaller, reaching that of the monomer and demonstrating that dimeric PpADK1 dissociates to monomer during a ternary complex formation to be active. We thus deduce that the dimer dissociation is likely rate-limiting for the catalysis of both moss enzymes, showing up to 10-fold lower kinase activities than all three maize mainly monomeric ADKs. How the substrate induces the dimer dissociation remains unclear.

In line with the above, soaking crystals of apo ZmADK2 with the substrates Ado or iPR or Ado and co-substrate analog AMP-PCP was unsuccessful due to the impossible rearrangement and closure of the substrate cavity within the dimer inside the crystals. Since the ATP site was accessible, we obtained a complex of ZmADK2 with AMP-PCP in the open conformation. The closed conformation was previously observed only in the structures where the substrate site was occupied, usually together with the ATP site, including human and Toxoplasma ADK with bound Ado (PDB 1BX4, Mathews et al., 1998; PDB 1LIK, 1LII and 1LIJ, Schumacher et al., 2000). It has also been observed for bound Ado with AMP-PNP (PDB 4N09) and AP5A in trypanosomal ADK structures (PDB 3OTX) (Kuettel et al., 2011).

In Arabidopsis, AtADK2 was shown to form a protein complex *in vivo* and *in vitro* with the sucrose non-fermenting 1 (SNF1)-related kinase abbreviated as SnRK1 (Mohannath et al., 2014). AMP prevents dephosphorylation and thus its inactivation. When various stresses deplete energy stored in ATP, increased levels of AMP maintain the activity of SnRK kinases, which turn off energy-consuming biosynthetic pathways and turn on alternative ATP-generating reactions (Halford and Hey, 2009). The N-terminal kinase domain of AtSnRK1 and its inactive K49R variant were sufficient to form a complex with AtADK2 and increase ADK activity up to seven times (Mohannath et al., 2014). It has been hypothesized that the AtADK2-AtSnRK1 complex may facilitate cellular stress responses and ADK may provide AMP to keep SnRK1 active. At the same time, the direct interaction with SnRK1 may lead to conformational changes in AtADK and increase ADK activity. For now, we can only speculate that AtSnRK1, for example, can prevent the dimerization of AtADK2 since its oligomeric state has never been reported. In addition, AtADK2 is also known to be inhibited by the geminivirus AL2 and L2 proteins, which also interact with and inactivate SnRK1 kinase (Wang et al., 2003). Here, we could suggest that one of the binding sites is partially blocked or the rotation of a small domain is affected.

### Substrate preferences among various ADKs

Substrate preferences have been analyzed in the past for ADKs from the human placenta (Andres and Fox, 1979), rabbit liver (Miller et al., 1979a), *T. gondii* (Iltzsch et al., 1995; Darling et al., 1999), *T. brucei* (Vodnala et al., 2008), and several other species. Substrate specificity screenings with rabbit and Toxoplasma ADKs showed Ado as the best substrate, followed by 7-deaza, 1-deaza and 3-deaza Ado derivatives. Poor activities of the latter two were explained by the missing H-bond interactions with the N1 and N3 atoms within the active site (Mathews et al., 1998). Many substituted nucleosides have been tested for antitubercular activity against mycobacterial ADK from *Mycobacterium tuberculosis* (MtADK, Long and Parker, 2006). For example, 2-methylAdo was a much better substrate and selective against human ADK (Long et al., 2003). It was shown that the difference in catalytic efficiency was three orders of magnitude in favor of MtADK when phosphorylating 2-methylAdo. The activities and *K*_m_ values measured in this work for the cytokinin ribosides iPR, ZR and BAPR with five plant ADKs (Table 1 and Table 2) align with previous publications for lupin, wheat and tobacco ADKs (Guranowski, 1979; Chen and Eckert, 1977; Kwade et al., 2005) as well as for two Arabidopsis ADKs (Moffatt et al., 2000).

Because of the remarkable conservation of the Ado pocket among plant (Figure 3I) and mammalian ADK sequences, it is reasonable to deduce that mammalian ADKs can naturally bind plant-derived cytokinin ribosides. Indeed, rabbit ADK has also been shown to phosphorylate cytokinin ribosides (Miller et al., 1979a). While *K*_m_ for Ado was 0.4 µM, those for iPR, ZR and BAPR were 7, 20 and 12 µM, respectively. The enzyme showed 69 % activity with iPR, 29% with BAPR and 6.3% with ZR compared to Ado. Our *in silico* docking analysis showed almost no differences in the binding of iPR to ZmADK3 and human ADK (Figure S8). Phe201 and Leu40 have to flip to leave more space for the side chain. However, the binding energies between Ado and iPR are comparable, suggesting similar binding modes. Moreover, previous analysis of rabbit ADK revealed that 2′-deoxyAdo and 3′-deoxyAdo derivatives had much higher *K*_m_ values of 575 and 254 µM than Ado itself (Miller et al., 1979a). Our data measured for ZmADK2 (Table 3) show a remarkable similarity in *K*_D_ and *K*_m_ values of ∼ 500 µM and 170 µM, respectively, for each ligand. This similarity can be attributed to preserving active site residues, particularly the aspartate residue (D17 in ZmADK2, D18 in human ADK), which binds the ribose substrate’s O2’ and O3’ hydroxyl groups.

Although ATP is the preferred phosphate donor, other molecules such as GTP, ITP, dATP and others have also been shown to be donors, although they are much weaker. Nevertheless, the reported *K*_m_ and affinity values are very inconsistent, ranging from low micromolar to low millimolar concentrations. For example, *K*_m_ values of 20 µM and 0.8 mM for ATP have been measured with rat ADK (Fisher and Newsholme, 1984; Yamada et al., 1980). The *K*_m_ values of 0.3-0.4 mM for ATP were measured with plant ADKs from lupin and Arabidopsis in the range of reported intracellular ATP levels (Guranowski, 1979; Moffatt et al., 2000).

Here, we used MST to determine *K*_D_ values, as high ATP concentrations inhibit the coupled reaction. Thus, the binding could be measured without interference from the Ado substrate. The calculated *K*_D_ value for ATP (in the presence of magnesium ions) with ZmADK2 was 2.2 ± 0.6 mM (Table 3). The affinity for ADP was even lower (*K*_D_ value of 10.3 ± 1.2 mM), which is logical since reaction products usually have lower affinities than substrates. The millimolar affinity for ATP was confirmed by Ado, showing the second binding event with a *K*_D_ value of ∼ 1.5 mM (Table 3 and Figure S6). Indeed, Ado has been described as a competitive inhibitor of ATP binding (Fisher and Newsholme, 1984) and Ado has been observed bound to the ATP site of human ADK (Mathews et al., 1998). Similarly, the AMP exhibited the first *K*_D_ value of ∼ 250 µM, corresponding to the product binding in the Ado pocket, and the second *K*_D_ value of ∼ 8 mM, corresponding to binding in the ATP pocket.

### Consequences of elevated ADK activity on metabolites and phenotype

It is well known that ADK activity maintains the rate of SAM-dependent transmethylation reactions in plants by reducing free Ado levels (Moffatt et al., 2002) and ADK activity has been observed to increase between 1.5- and 2.8-fold in response to methyl demand (Weretilnyk et al., 2001). Here, we focused on the relevance of ADK activity in nitrogen salvage, which was analyzed in the lines overexpressing *ZmADK1/2/3*. Lack of nitrogen induces the degradation of RNA (Kraft et al., 2008), which enables plants to recycle the reduced nitrogen. We have recently shown evidence of accelerated purine and pyrimidine catabolism towards amino acids and polyamines in *pOpON::ZmNRH* transgenics (Ľuptáková et al., 2024). Although NRHs catalyze the opposite reaction from Ado to adenine, it can be converted to AMP by APTs. In addition, NRHs also act downstream of ADK in the purine catabolic pathway by hydrolyzing xanthosine to xanthine, which has been shown to act as a nitrogen source (Melino et al., 2018). Levels of uridine, xanthosine, inosine, guanosine or cytokinin riboside iPR were significantly decreased while the Ado levels were compensated. Here, in *pOpON::ZmADK* transgenics, Ado, cytokinin ribosides and xanthosine were among the significantly decreased ribosides. It is not surprising that levels of Ado, iPR, *t*ZR or *c*ZR were increased in *ADK*-silenced Arabidopsis lines (Schoor et al., 2011), while AMP levels were not decreased but similar or even elevated.

Abiotic stress and nitrogen-limiting conditions result in increased chlorophyll bleaching, RNA breakdown and growth retardation. Overexpression of ureide permease, another enzyme in the purine degradation pathway, increased chlorophyll content, tiller number, shoot, and root biomass in nitrogen-starved rice (Redillas et al., 2019). Root elongation in *ZmNRH2b/3* overexpressor lines was about 50% faster under nitrogen starvation than WT; the rosette area and green leaf index were larger (Ľuptáková et al., 2024). Here, we observed a statistically higher number of lateral roots and larger root area in nitrogen-starved *ZmADK* overexpressor lines, showing benefits similar to those of *ZmNRH* overexpressors. On the other hand, no phenotypic effects were observed on the shoot concerning plant size, rosette area or seed number. This contrasts *ADK*-silenced Arabidopsis lines (Moffatt et al., 2002), which showed distinct phenotypes, including reduced root and stem elongation, bushier rosettes, and stamen and silique development defects. However, the reason was attributed to the inhibition of the SAM cycle and reduced methylation reactions but not directly to the purine catabolism pathway. We can summarize that the effect of *ADK* overexpression on phenotype is milder than that of *NRH* because ADK stands at the beginning of the purine catabolism and released AMP can be diverted to the ATP synthesis and the interconversion to other purine metabolites before its conversion to xanthine.

## SUPPLEMENTARY DATA

**Table S1.** Data collection and refinement statistics of maize ADKs.

**Table S2.** Primer pairs and probes used for RT-qPCR determination.

**Table S3.** Results of docking calculations into both sites in selected ADK isoforms.

**Table S4.** Transcript abundance of three *ADK* genes in maize tissues.

**Table S5.** Transcript abundance of three *ADK* genes in moss.

**Table S6.** Nucleoside levels in *A. thaliana ZmADK* overexpressors.

**Table S7.** Total root area of *pOpOn::ZmADK* transgenic lines in nitrogen-varying conditions.

**Table S8.** Change of leaf area in *pOpOn::ZmADK* transgenic lines in nitrogen-varying conditions.

**Table S9.** Levels change among selected amino acids in *pOpOn::ZmADK* transgenic lines in the absence of nitrogen.

**Figure S1.** Thermal stability, molecular and kinetic properties focused on ZmADK2.

**Figure S2.** Reaction scheme for the synthesis of two purine ribosides.

**Figure S3.** NMR spectra of *N^6^*-methylAdo.

**Figure S4.** NMR spectra of *N^6^*, *N^6^*-dimethylAdo.

**Figure S5.** NMR spectra of *N^6^*-isopropylAdo.

**Figure S6.** NMR spectra of *N^6^*-isobutylAdo.

**Figure S7.** Sequence alignment of the selected plant ADKs and human ADK.

**Figure S8.** *In silico* docking of cytokinin riboside in the active site of ADK.

**Figure S9.** Comparison between the two known ADK dimers.

## ACKNOWLEDGMENTS

We acknowledge SOLEIL for providing of synchrotron radiation facilities in using PROXIMA 1 and 2 beamlines (proposals ID 20170872, 20191181 and 20210831). We would like to thank Mgr. Anna Krnáčová for HPLC-MS analysis and Pharm.Dr. Jitka Široká, Ph.D. for measurement of HRMS data. We acknowledge CF Biomolecular Interactions and Crystallography of CIISB, Instruct-CZ Centre, for the assistance of AUC measurements, supported by MEYS CR (LM2023042) and European Reginoval Development Fund-Project “UP CIISB” (No. CZ.02.1.01/0.0/0.0/18_046/0015974).

## AUTHOR CONTRIBUTIONS

DK, MS and SM designed the research. DJK, MK, JB, RK and KvS prepared enzyme variants and their mutants, as well as measured enzyme properties with kinetics and real-time qPCR; VM synthesized new ligands; DJK prepared transgenic lines; JFH and DJK performed the phenotyping analysis; JF, MS and ON were involved in measurements and analysis of purine metabolites *in vivo*; AV, MLB, SP, DK and SM performed the crystallographic study and analyzed the crystal structures; DK and SM wrote the paper. All authors reviewed the results and approved the final version of the manuscript submitted for publication.

## DATA AVAILABILITY

The atomic coordinates and structure factors have been deposited in the Protein Data Bank (www.wwpdb.org) under accession codes 8RF7 for ZmADK2 apoenzyme, 8RGJ for ZmADK2 complex with AMP-PCP and 8RPA for the ZmADK3 complex with AP5A.

