## Supplementary material for "The monomer/dimer switch modulates the activity of plant adenosine kinase": Table S1

**Table S1**. **Data collection and refinement statistics.**

| Enzyme | **ZmADK2** | **ZmADK2** | **ZmADK3** | **PpADK1** |
|---|---|---|---|---|
| Observed ligand | - | AMP-PCP | AP5A | Ado + ADP |
| PDB code | 8RF7 | 8RGJ | 8RPA | 9FW6 |
| Space group | P4212 | P4212 | I212121 | P212121 |
| Asymmetric unit | 1 dimer | 1 dimer | 1 monomer | 2 monomers |
| Unit cell (Å) | | | | |
| a | 136.22 | 135.85 | 51.0 | 50.59 |
| b | 136.22 | 135.85 | 117.5 | 93.08 |
| c | 78.23 | 78.63 | 166.0 | 131.67 |
| α (°) | 90.0 | 90.0 | 90.0 | 90.0 |
| β (°) | 90.0 | 90.0 | 90.0 | 90.0 |
| γ (°) | 90.0 | 90.0 | 90.0 | 90.0 |
| Resolution (Å) | 96.3 – 2.05 (2.17 – 2.05) | 96.1 – 2.36 (2.51 – 2.36) | 48.0 – 2.26 (2.54 – 2.26) | 76.0 – 1.73 (1.93 – 1.73) |
| Resolution limits by Staraniso (Å) | 2.05, 2.05, 3.02 | 2.36, 2.36, 3.31 | 3.11, 2.52, 2.26 | 2.48, 1.73, 2.05 |
| Observed reflections | 800280 (41504) | 535990 (28034) | 162656 (6984) | 442676 (23441) |
| Unique reflections | 30369 (1520) | 20930 (1046) | 14056 (703) | 37232 (1862) |
| Completeness (spherical) (%) | 65.2 (19.2) | 67.7 (20.8) | 58.8 (10.2) | 56.7 (10.1) |
| Completeness (ellipsoidal) (%) | 94.9 (80.4) | 93.7 (78.9) | 89.8 (68.6) | 93.1 (82.2) |
| $I/\sigma(I)$ | 15.6 (1.5) | 12.4 (1.5) | 8.5 (1.4) | 6.1 (1.7) |
| $R_{sym}$ (%) | 20.4 (287.9) | 31.4 (285.6) | 26.3 (195.5) | 31.5 (233.5) |
| $R_{pim}$ (%) | 4.0 (55.9) | 6.3 (55.9) | 7.7 (57.1) | 9.5 (67.8) |
| $CC_{1/2}$ | 99.9 (40.5) | 99.7 (51.1) | 99.4 (65.2) | 98.4 (61.0) |
| Amino acid residues | 667 | 671 | 341 | 670 |
| Water molecules | 174 | 147 | 99 | 320 |
| Ligand molecules | - | 2 | 1 | 4 |
| $R_{cryst}$ (%)[c] | 23.4 | 26.0 | 20.8 | 22.0 |
| $R_{free}$ (%) | 26.1 | 27.8 | 25.7 | 25.5 |
| RMSD bond lengths (Å) | 0.08 | 0.007 | 0.007 | 0.009 |
| RMSD bond angles (°) | 0.97 | 0.88 | 0.88 | 1.00 |
| Mean B value (Å$^2$): | | | | |
| Overall | 48.9 | 56.2 | 31.2 | 30.3 |
| protein chain A/B | 49.9/48.5 | 55.5/55.8 | 31.6 | 33.1/27.8 |
| solvent molecules | 49.6 | 54.3 | 21.4 | 25.9 |
| ligand A/B | -/- | 74.6/84.2 (AMP-PCP) | 29.2 (AP5A) | 34.4/31.4 (ADP) 21.5/17.5 (Ado) |
| Ramachandran statistics (%)[d]: | | | | |
| Favored | 99.24 | 95.92 | 97.1 | 97.90 |
| Outliers | 0.0 | 0.3 | 0.0 | 0.0 |
| Clash score (PR)[d] | 1.27 | 1.16 | 0.75 | 0.39 |
| MolProbity score (PR)[d] | 0.85 | 1.11 | 1.10 | 0.70 |

[a] Numbers in parentheses represent values in the highest resolution shell.
[b] $CC_{1/2}$ represents a percentage of correlation between intensities from a random half-dataset.
[c] The 5% test set.
[d] Generated with MolProbity.
