## Supplementary material for "The monomer/dimer switch modulates the activity of plant adenosine kinase": Table S2

**Table S2. Primer pairs and probes used for RT-qPCR determination.**

| | Primer pairs and probes for RT-qPCR |
|---|---|
| *ZmADK1* | 5´- AGGTCCTCCCGTATGCTGACT -3´,<br>5´- CTGTCTCCCATCCTCGAACTTT -3´,<br>5´-FAM- CATCTTCGGCAATGAAACCGAGGC -TAM -3´ |
| *ZmADK2* | 5´- AGGTGAAAACGTTCCCTGTGA -3´,<br>5´- CAATGCCCTTTCCTAGAACCAA -3´,<br>5´-FAM- ATGCTTTCGTTGGAGGCTTCCTCTCAC -TAM -3´ |
| *ZmADK3* | 5´- TCCCCGGATTCTATTCAACTTG -3´,<br>5´- GGAGCAGAAAGGTTCATCATAAACA -3´,<br>5´-FAM- TGCTGAGCATGCCGCTGCAAC -TAM -3´ |
| *PpADK1* | 5´- CGATACGATGTGGACGAGGAT -3´,<br>5´- CCAAGGACCTCTCTCCCTTCA -3´,<br>5´-FAM- TTCCCACTGGAACATGCGGCG -TAM -3´ |
| *PpADK2* | 5´- GGCTGCGTTGCCCAAA -3´,<br>5´- GGGATCAGTTCCCTGAGTGATAA -3´,<br>5´-FAM- CTAGCGGCACTCACAAGCGTGTTGC -TAM -3´ |
| *PpADK3* | 5´- TCATTTGTTTGTGAGCGTTTTAAAG -3´,<br>5´- GCCTCATTCCCAAACATGTAGTC -3´,<br>5´-FAM- TCCTCTCATGGCAGCCTTCCCATATG -TAM -3´ |
| *AtNRH1* | 5´-TCTAGATGAGAAGGTCGAAGAATATCC-3´,<br>5´-GGCTAATGCCAGGTTGGTTAGAG-3´,<br>5´-FAM-TGAAGTCACCATTCTCGCCCTCGG-TAM-3´ |
| *AtNRH2* | 5´-CCTCACTACTATCTTTGGAAACGTGTA-3´,<br>5´-CGCAACCTCCAACAAATGC-3´,<br>5´-FAM- CCACTCTCGCCACTCGAAACGCC-TAM-3´ |
| *AtADK1* | 5´-GTCGCTGAGGATGGAAAAGTG-3´,<br>5´-GCACCGTTGGTGTCAACAAG-3´,<br>5´-FAM- AGAAGTACCCAGTCATCCCTCTCCCCAA-TAM-3´ |
| *AtADK2* | 5´-TCGAACAAATAGCCATCAAGATTTC-3´,<br>5´-CAGCGCCCTGTGTAATCACA-3´,<br>5´-FAM- CCCAAGGCCACAGGAACATACAAGAGG-TAM-3´ |
| *AtAPT1* | 5´-CTCAAACCACCGTTCAACCA-3´,<br>5´-CGGCGTCAGATTTGCAAAC-3´,<br>5´-FAM- CCTCTTCTCCTCCGCCGGGTCTC-TAM-3´ |
| *AtAPT3* | 5´-GGCTCGTGGTTTCCTATTCG-3´,<br>5´-GTTTGCGCAGAGGAACAAATTT3´,<br>5´-FAM- TCCACCGATCGCGCTAGCCATT-TAM-3´ |
| *AtLOG7* | 5´-TATGCATCAAAGGAAAGCTGAAAT-3´,<br>5´-AACGTACCATACCCACCAGGAA-3´,<br>5´-FAM- CTCGCCAAGCCGACGCATTCA-TAM-3´ |
| *AtLOG8* | 5´-CAGCGAAGCAGATTCAGAAAAA-3´,<br>5´-GGCAGCATCACTGAAAACTTCTC-3´,<br>5´-FAM- TCTTTTGCGGAAGCCACTCTGGTCA-TAM-3´ |
