## Supplementary material for "The monomer/dimer switch modulates the activity of plant adenosine kinase": Table S3

**Table S3. Results of docking calculations into both sites in selected ADK isoforms.** Docked using FLARE software (www.cresset-group.com/software/flare-docking/). The rank score is designed to provide the best score for the correct (experimentally observed) ligand pose; ΔG represents an accurate estimation of the free energy of protein-ligand binding for a given protein-ligand complex.

| | **Ligand** | ***ZmADK2* *(8RGJ, opened)*** | | ***ZmADK3* *(8RPA; closed)*** | | ***PpADK1* *(9FW6; closed)*** | | ***HsADK* *(1BX4; closed)*** | |
|---|---|---|---|---|---|---|---|---|---|
| | | Rank Score | Δ*G score* | Rank Score | Δ*G* | Rank Score | Δ*G* | Rank Score | Δ*G* |
| | | (kcal mol$^{-1}$) | | | | | | | |
| *Ado-site* | *Ado* | -4.6 | -4.4 | -10.3 | -7.5 | -10.1 | -8.5 | -10.4 | -8.0 |
| | *AMP* | -4.7 | -5.0 | -9.5 | -7.8 | -10.6 | -8.6 | -7.4 | -7.2 |
| | *iPR* | -5.2 | -5.5 | -8.6 | -8.6 | -9.8 | -9.8 | -9.3 | -9.0 |
| | *BAPR* | -5.9 | -4.9 | -9.2 | -8.3 | -10.0 | -9.3 | -9.6 | -8.1 |
| *ATP-site* | *Ado* | n. d. | n. d. | -5.0 | -5.3 | -6.6 | -5.9 | -7.0 | -6.4 |
| | *AMP* | -3.9 | -4.4 | -3.2 | -4.3 | -5.7 | -5.7 | -6.4 | -5.8 |
| | *ADP* | -3.3 | -4.4 | -3.3 | -4.7 | -6.9 | -6.1 | -6.0 | -5.6 |
| | *ATP* | -3.3 | -4.3 | -3.4 | -3.6 | -7.3 | -6.2 | -5.2 | -5.5 |
