## Supplementary material for "The monomer/dimer switch modulates the activity of plant adenosine kinase": Table S4

**Table S4. Transcript abundance of three *ADK* genes in maize tissues.** Abundance is the number of transcripts per ng of total RNA amplified by qPCR. RNA from four biological replicates was transcribed in two independent reactions and PCR was performed in duplicate. Mean values ± standard deviations are shown. DAG, days after germination; DAP, days after pollination; DBP, days before pollination; M, months.

| | **Gene (transcripts ng$^{-1}$ of total RNA)** | | |
|---|---|---|---|
| **Sample** | ***ZmADK1*** | ***ZmADK2*** | ***ZmADK3*** |
| embryo | 106.2 ± 11.3 | 1160.6 ± 129.4 | 134.9 ± 11 |
| stem (9 DAG) | 187.8 ± 14.4 | 336.1 ± 16.8 | 9.9 ± 1.2 |
| leaf (9 DAG) | 256.2 ± 26 | 277.6 ± 27.7 | 71.3 ± 4.3 |
| root (9 DAG) | 50 ± 8.3 | 293.8 ± 23.6 | 12.1 ± 2.7 |
| stem (3 M) | 172.1 ± 17.4 | 883.8 ± 78.9 | 94.1 ± 5.7 |
| leaves (3 M) | 130.3 ±10.1 | 1145.4 ± 113.2 | 141.3 ± 31 |
| main root (3 M) | 88.5 ± 4.4 | 697.7 ± 20 | 28.9 ± 2.5 |
| tassels (5 DBP) | 98.6 ± 3.6 | 731.6 ± 120.4 | 28.9 ± 2.9 |
| tassels (0 DAP) | 555 ± 51.8 | 2923.6 ± 390.7 | 227.8 ± 50 |
| silks (0 DAP) | 163.9 ± 2.2 | 572.3 ± 103.9 | 53.2 ± 2.6 |
| silks (3 DAP) | 80 ± 2.3 | 2773.8 ± 314.6 | 337.1 ± 85.7 |
| silks (6 DAP) | 109.5 ± 3.2 | 2373.4 ± 356.9 | 1003.2 ± 141.1 |
| kernel (3 DBP) | 351.5 ± 20.9 | 649.1 ± 68.8 | 121.8 ± 14.8 |
| kernel (3 DAP) | 801.5 ± 23.7 | 1263.1 ± 177 | 504.9 ± 115.7 |
| kernel (9 DAP) | 1353.1 ± 417.1 | 1345 ± 157.5 | 313.4 ± 48.9 |
| kernel (15 DAP) | 7634.4 ± 425.3 | 10611 ± 1975.6 | 2371.2 ± 139.7 |
| kernel (20 DAP) | 1869.5 ± 163.5 | 3522.5 ± 514.4 | 1636.3 ± 150.4 |
| ***Stem*** | | | |
| 3 DAG | 140.5 ± 10.5 | 304.6 ± 30.2 | 10.2 ± 0.7 |
| 5 DAG | 94.7 ± 4.9 | 243.4 ± 13.9 | 5.7 ± 1.6 |
| 7 DAG | 75.1 ± 10.3 | 215.5 ± 33.0 | 3.8 ± 0.9 |
| 9 DAG | 205.9 ± 34.4 | 385.1 ± 35.1 | 5.7 ± 0.5 |
| 11 DAG | 187.8 ± 16.8 | 336.1 ± 58.1 | 9.9 ± 1.1 |
| 13 DAG | 202.1 ± 23.3 | 279.4 ± 9.0 | 13.2 ± 3.1 |
| ***Leaves*** | | | |
| 3 DAG | 28.3 ± 4.5 | 153.6 ± 20.8 | 257.8 ± 18.9 |
| 5 DAG | 11.4 ± 1.4 | 78.5 ± 6.7 | 157.3 ± 25.1 |
| 7 DAG | 10.5 ± 0.9 | 163.1 ± 5.3 | 120.3 ± 19.2 |
| 9 DAG | 40.2 ± 5.8 | 310.8 ± 32.1 | 249.6 ± 39.9 |
| 11 DAG | 50.2 ± 4.3 | 293.8 ± 19.8 | 459.8 ± 73.1 |
| 13 DAG | 62.5 ± 3.2 | 286.5 ± 37.7 | 140.8 ± 22.5 |
| ***Roots*** | | | |
| 3 DAG | 71.5 ± 11.5 | 298.2 ± 17.2 | 23.7 ± 3.7 |
| 5 DAG | 118.8 ± 15.6 | 515.6 ± 22.9 | 74.1 ± 11.8 |
| 7 DAG | 135.5 ± 15.5 | 370.3 ± 25.6 | 78.9 ± 12.6 |
| 9 DAG | 138.2 ± 9.0 | 216.1 ± 18.7 | 32.2 ± 1.9 |
| 11 DAG | 365.3 ± 56.6 | 277.6 ± 14.5 | 44.1 ± 2.2 |
| 13 DAG | 144.3 ± 18.9 | 383.7 ± 31.6 | 36.9 ± 7.3 |
| ***Leaves at 9 DAG*** | | | |
| No treatment | 156.2 ± 26.2 | 277.6 ± 27.7 | 23.6 ± 1.1 |
| + 200 mM NaCl | 215 ± 15.1 | 175.9 ± 11.8 | 71.7 ± 5.9 |
| - Nitrogen | 76.7 ± 8.4 | 281.7 ± 24.6 | 68.6 ± 8.2 |
| + 1 µM *t*Z | 109.6 ± 13.2 | 254.3 ± 33.0 | 49.8 ± 5.7 |
| ***Roots at 9 DAG*** | | | |
| No treatment | 120 ± 8.3 | 293.8 ± 23.6 | 32.1 ± 2.7 |
| + 200 mM NaCl | 220.6 ± 15.4 | 227.5 ± 20.8 | 26.4 ± 1.8 |
| - Nitrogen | 46.3 ± 4.7 | 84 ± 9.6 | 28.5 ± 3.1 |
| + 1 µM *t*Z | 89.3 ± 10.7 | 141.6 ± 12.3 | 49.3 ± 5.9 |
