## Supplementary material for "The monomer/dimer switch modulates the activity of plant adenosine kinase": Table S5

**Table S5. Transcript abundance of three *ADK* genes in moss.** Abundance is the number of transcripts per ng of total RNA amplified by qPCR. RNA from three technical replicates. Mean values ± standard deviations are shown. N.D., not detected

| | Gene (transcripts $ng^{-1}$ of total RNA) | | |
|---|---|---|---|
| **Sample** | ***PpADK1*** | ***PpADK2*** | ***PpADK3*** |
| Control | 13853 ± 266 | 78 ± 3 | N.D. |
| + 200 mM NaCl | 22342 ± 826 | 120 ± 3 | N.D. |
| - Nitrogen | 14011 ± 579 | 64 ± 6 | N.D. |
| + Mannitol | 30570 ± 832 | 156 ± 10 | N.D. |
| + 10 µM ABA | 18401 ± 516 | 170 ± 10 | N.D. |
| + 10 µM cytokinin (BAP) | 15309 ± 181 | 87 ± 6 | N.D. |
| + 10 µM auxin (2,4-D) | 8482 ± 314 | 61 ± 1 | 3 ± 0 |
