## Supplementary material for "The monomer/dimer switch modulates the activity of plant adenosine kinase": Table S6

**Table S6. Nucleoside levels in *A. thaliana ZmADK* overexpressors.** Values are given in nmol $g^{-1}$ for adenine ribosides and in pmol $g^{-1}$ for cytokinin ribosides. Data were measured after dexamethasone induction at 72 hours. Asterisks indicate statistically significant differences in treated lines versus the controls (non-treated plants) in a paired Student's t-test (t-test; *, **, and *** correspond to P-values of 0.05 > p > 0.01, 0.01 > p > 0.001, and p < 0.001, respectively). Levels of IMP, XMP and GMP were below the limit of detection.

| | **Metabolites (pmol $g^{-1}$ FW)** | | | | |
|---|---|---|---|---|---|
| | Ade | Ado | Guanosine | Xanthosine | Inosine |
| WT | 1910.7 ± 305.9 | 8.4 ± 1.2 | 194.4 ± 14.7 | 361.5 ± 13.9 | 52.9 ± 2.5 |
| *ZmADK1 #10-6* | 1750.1 ± 145.3 | 5.4 ± 0.5* | 138.3 ± 7.3** | 364.7 ± 10.1 | 82.4 ± 1.9 |
| *ZmADK1 #11-4* | 1515.4 ± 164.3 | 4.6 ± 1.2** | 155.4 ± 12.7* | 238.4 ± 4.9*** | 113.1 ± 6.5** |
| *ZmADK2 #2-4* | 1427.3 ± 49.8* | 4.8 ± 1.1** | 131.6 ± 9.7** | 248.5 ± 16.1*** | 133.9 ± 2.3*** |
| *ZmADK3 #3-3* | 1446.5 ± 78.4 | 3.4 ± 1.0** | 154.8 ± 16.7* | 184.9 ± 7.5*** | 77.3 ± 6.2 |
| *ZmADK3 #11-14* | 1801.0 ± 31.0 | 4.7 ± 1.2** | 154.1 ± 9.7* | 221.6 ± 9.9*** | 51.2 ± 2.3 |

| | Cytokinin bases | Cytokinin ribosides | Cytokinin monophosphates | iP | iPR | iPRMP |
|---|---|---|---|---|---|---|
| WT | 0.51 ± 0.06 | 4.33 ± 0.25 | 14.08 ± 0.13 | 0.32 ± 0.05 | 1.24 ± 0.07 | 7.89 ± 0.32 |
| *ZmADK1 #10-6* | 0.50 ± 0.05 | 2.99 ± 0.13** | 12.78 ± 0.30** | 0.32 ± 0.04 | 0.94 ± 0.05** | 6.67 ± 0.32** |
| *ZmADK1 #11-4* | 0.59 ± 0.05 | 2.08 ± 0.27*** | 13.40 ± 0.33* | 0.37 ± 0.06 | 0.75 ± 0.16** | 8.54 ± 0.13* |
| *ZmADK2 #2-4* | 0.59 ± 0.03 | 2.24 ± 0.28*** | 12.40 ± 0.28*** | 0.36 ± 0.05 | 0.74 ± 0.14** | 7.70 ± 0.23 |
| *ZmADK3 #3-3* | 0.49 ± 0.04 | 2.16 ± 0.05*** | 12.83 ± 0.65* | 0.29 ± 0.03 | 0.85 ± 0.09** | 8.05 ± 0.32 |
| *ZmADK3 #11-14* | 0.49 ± 0.03 | 2.62 ± 0.10*** | 13.89 ± 0.14 | 0.30 ± 0.03 | 0.90 ± 0.14* | 8.02 ± 0.10 |

| | *c*Z | *c*ZR | *c*ZRMP | *t*ZR | DHZR |
|---|---|---|---|---|---|
| WT | 0.15 ± 0.01 | 1.78 ± 0.16 | 5.37 ± 0.21 | 1.20 ± 0.16 | 0.11 ± 0.01 |
| *ZmADK1 #10-6* | 0.15 ± 0.01 | 1.27 ± 0.10** | 5.62 ± 0.31 | 0.71 ± 0.09** | 0.08 ± 0.01* |
| *ZmADK1 #11-4* | 0.16 ± 0.01 | 0.58 ± 0.07*** | 4.35 ± 0.38* | 0.70 ± 0.04** | 0.05 ± 0.01** |
| *ZmADK2 #2-4* | 0.17 ± 0.01 | 0.73 ± 0.15** | 4.17 ± 0.14** | 0.72 ± 0.05** | 0.04 ± 0.01*** |
| *ZmADK3 #3-3* | 0.15 ± 0.01 | 0.69 ± 0.18** | 4.26 ± 0.31** | 0.58 ± 0.05** | 0.03 ± 0.01*** |
| *ZmADK3 #11-14* | 0.17 ± 0.01 | 1.06 ± 0.06** | 5.34 ± 0.24 | 0.61 ± 0.04** | 0.07 ± 0.02 |
