## Supplementary material for "The monomer/dimer switch modulates the activity of plant adenosine kinase": Table S8

**Table S8. Change of leaf area in *pOpOn::ZmADK* transgenic lines in nitrogen-varying conditions.** At least 24 seedlings (biological replicates) per and growth condition were included. Median values, mean and standard error values are shown. Asterisks indicate statistically significant differences in transgenic lines versus the WT controls in a paired Student's t-test (t-test; *, **, and *** correspond to P-values of 0.05 > p > 0.01, 0.01 > p > 0.001, and p < 0.001).

| ***No nitrogen*** | *Stage* | *Median* | *Mean* | *St. error* | *p-value* |
|---|---|---|---|---|---|
| *WT* | 6 DAG | 307 | 330.8 | 26.4 | |
| *ZmADK1 #11-4* | 6 DAG | 348 | 363.3 | 27.8 | - |
| *ZmADK3 #3-3* | 6 DAG | 316 | 302.3 | 33.1 | - |
| *ZmADK3 #11-14* | 6 DAG | 285 | 294.4 | 21.8 | - |
| *WT* | 8 DAG | 480 | 518.5 | 43.5 | |
| *ZmADK1 #11-4* | 8 DAG | 534 | 554.0 | 65.4 | - |
| *ZmADK3 #3-3* | 8 DAG | 531 | 574.5 | 44.7 | - |
| *ZmADK3 #11-14* | 8 DAG | 467 | 468.2 | 29.7 | - |
| *WT* | 10 DAG | 638 | 646.0 | 57.2 | |
| *ZmADK1 #11-4* | 10 DAG | 610 | 627.3 | 87.6 | - |
| *ZmADK3 #3-3* | 10 DAG | 710 | 668.0 | 97.4 | - |
| *ZmADK3 #11-14* | 10 DAG | 600 | 643.5 | 72.4 | - |
| *WT* | 14 DAG | 944 | 915.7 | 82.6 | |
| *ZmADK1 #11-4* | 14 DAG | 970 | 1054.2 | 90.1 | * |
| *ZmADK3 #3-3* | 14 DAG | 916 | 971.3 | 99.6 | - |
| *ZmADK3 #11-14* | 14 DAG | 920 | 914.3 | 50.9 | - |
| ***Normal nitrogen*** | | | | | |
| *WT* | 6 DAG | 3940 | 3910.5 | 240.4 | |
| *ZmADK1 #11-4* | 6 DAG | 6147 | 5644.3 | 336.7 | *** |
| *ZmADK3 #3-3* | 6 DAG | 5544 | 5361.0 | 314.8 | ** |
| *ZmADK3 #11-14* | 6 DAG | 3145 | 3518.0 | 402.4 | - |
| *WT* | 8 DAG | 8480 | 8393.7 | 544.9 | |
| *ZmADK1 #11-4* | 8 DAG | 12239 | 11251.9 | 689.5 | ** |
| *ZmADK3 #3-3* | 8 DAG | 11132 | 11067.7 | 623.5 | ** |
| *ZmADK3 #11-14* | 8 DAG | 8555 | 8253.5 | 814.6 | - |
| *WT* | 10 DAG | 11028 | 11402.0 | 759.5 | |
| *ZmADK1 #11-4* | 10 DAG | 18318 | 17168.5 | 881.4 | *** |
| *ZmADK3 #3-3* | 10 DAG | 15492 | 15389.6 | 894.3 | ** |
| *ZmADK3 #11-14* | 10 DAG | 11586 | 12115.8 | 1450.2 | - |
| *WT* | 14 DAG | 28241 | 27831.4 | 2098.1 | |
| *ZmADK1 #11-4* | 14 DAG | 28703 | 28835.3 | 1668.4 | - |
| *ZmADK3 #3-3* | 14 DAG | 25451 | 27827.6 | 2034.2 | - |
| *ZmADK3 #11-14* | 14 DAG | 25399 | 27175.2 | 3194.5 | - |
| ***Double nitrogen*** | | | | | |
| *WT* | 6 DAG | 3124 | 3315.4 | 259.4 | |
| *ZmADK1 #11-4* | 6 DAG | 4884 | 4544.3 | 315.1 | ** |
| *ZmADK3 #3-3* | 6 DAG | 4140 | 4878.8 | 418.6 | ** |
| *ZmADK3 #11-14* | 6 DAG | 3989 | 3776.8 | 304.7 | - |
| *WT* | 8 DAG | 8247 | 7452.1 | 587.1 | |
| *ZmADK1 #11-4* | 8 DAG | 10156 | 9330.8 | 692.2 | * |
| *ZmADK3 #3-3* | 8 DAG | 9180 | 9520.3 | 775.0 | * |
| *ZmADK3 #11-14* | 8 DAG | 7480 | 6891.2 | 549.9 | - |
| *WT* | 10 DAG | 9982 | 10176.9 | 791.8 | |
| *ZmADK1 #11-4* | 10 DAG | 14209 | 13534.8 | 872.4 | ** |
| *ZmADK3 #3-3* | 10 DAG | 13368 | 12810.3 | 900.7 | * |
| *ZmADK3 #11-14* | 10 DAG | 11332 | 10355.1 | 797.1 | - |
| *WT* | 14 DAG | 15286 | 16580.6 | 1675.2 | |
| *ZmADK1 #11-4* | 14 DAG | 23796 | 23057.8 | 1383.9 | ** |
| *ZmADK2 #2-4* | 14 DAG | 23537 | 23816.0 | 1531.6 | ** |
| *ZmADK3 #3-3* | 14 DAG | 21076 | 20896.9 | 1631.2 | * |
| *ZmADK3 #11-14* | 14 DAG | 19198 | 18943.2 | 1585.0 | - |
