## Supplementary material for "The monomer/dimer switch modulates the activity of plant adenosine kinase": Table S9

**Table S9. Levels change among selected amino acids in pOpOn::ZmADK transgenic lines in the absence of nitrogen.** Values are given in nmol $g^{-1}$ DW with standard errors. Data were measured after dexamethasone induction for 48 hours in technical triplicates. Asterisks indicate statistically significant differences in transgenic lines versus the WT controls in a paired Student's t-test (t-test; *, **, and *** correspond to P-values of 0.05 > p > 0.01, 0.01 > p > 0.001, and p < 0.001).

| | **Metabolites (nmol $g^{-1}$ DW)** | | | | | |
|---|---|---|---|---|---|---|
| | Arg | Citrulline | Gln | Glu | Asn | Asp |
| WT | 363.0 ± 53.5 | 107.2 ± 15.6 | 48.2 ± 23.8 | 62.1 ± 8.9 | 151.2 ± 7.1 | 81.1 ± 16.5 |
| *ZmADK1 #10-6* | 470.5 ± 18.1* | 116.5 ± 9.5 | 103.2 ± 3.1* | 72.3 ± 29.1 | 179.5 ± 6.2* | 91.4 ± 12.0 |
| *ZmADK1 #11-4* | 552.4 ± 104.0* | 128.9 ± 18.8 | 121.4 ± 11.8** | 81.5 ± 33.2 | 188.3 ± 2.4** | 102.7 ± 12.1 |
| *ZmADK2 #2-4* | 529.3 ± 13.3** | 125.2 ± 8.7 | 145.9 ± 25.8** | 73.5 ± 28.0 | 161.5 ± 7.3 | 86.8 ± 20.9 |
| *ZmADK3 #3-3* | 516.5 ± 90.2 | 125.3 ± 20.4 | 95.8 ± 14.3* | 86.9 ± 23.0 | 169.1 ± 16.5 | 99.5 ± 10.8 |
| *ZmADK3 #11-14* | 564.9 ± 62.7* | 154.9 ± 20.1* | 99.7 ± 6.0* | 84.7 ± 9.7* | 164.7 ± 21.8 | 101.6 ± 6.0 |
