## Supplementary material for "The monomer/dimer switch modulates the activity of plant adenosine kinase": Figure S1

**Figure S1. Thermal stability, molecular and kinetic properties of ZmADK2. (A)** Thermal stability of five plant ADKs measured by nanoDSF. The curves for apoforms (in 150 mM HEPES pH 7.2, 100 mM NaCl, 10 mM $MgCl_2$) and ATP complexes at 4 mM concentrations are shown. **(B)** Gel filtration of highly concentrated ZmADK2 (at 30 mg ml-1) on Superdex S200 column. The chromatogram shows the existence of a small peak corresponding to the double molecular weight of the monomer. **(C)** Binding curves of Ado derivatives measured by MST in 50 mM HEPES buffer pH 7.5, 1 mM $MgCl_2$ and 0.1% Tween. **(D)** Saturation curves of Ado derivatives measured in a coupled reaction with PK and LDH at 30°C in 50 mM Tris-HCl buffer pH 7.5 using 10 mM ATP.

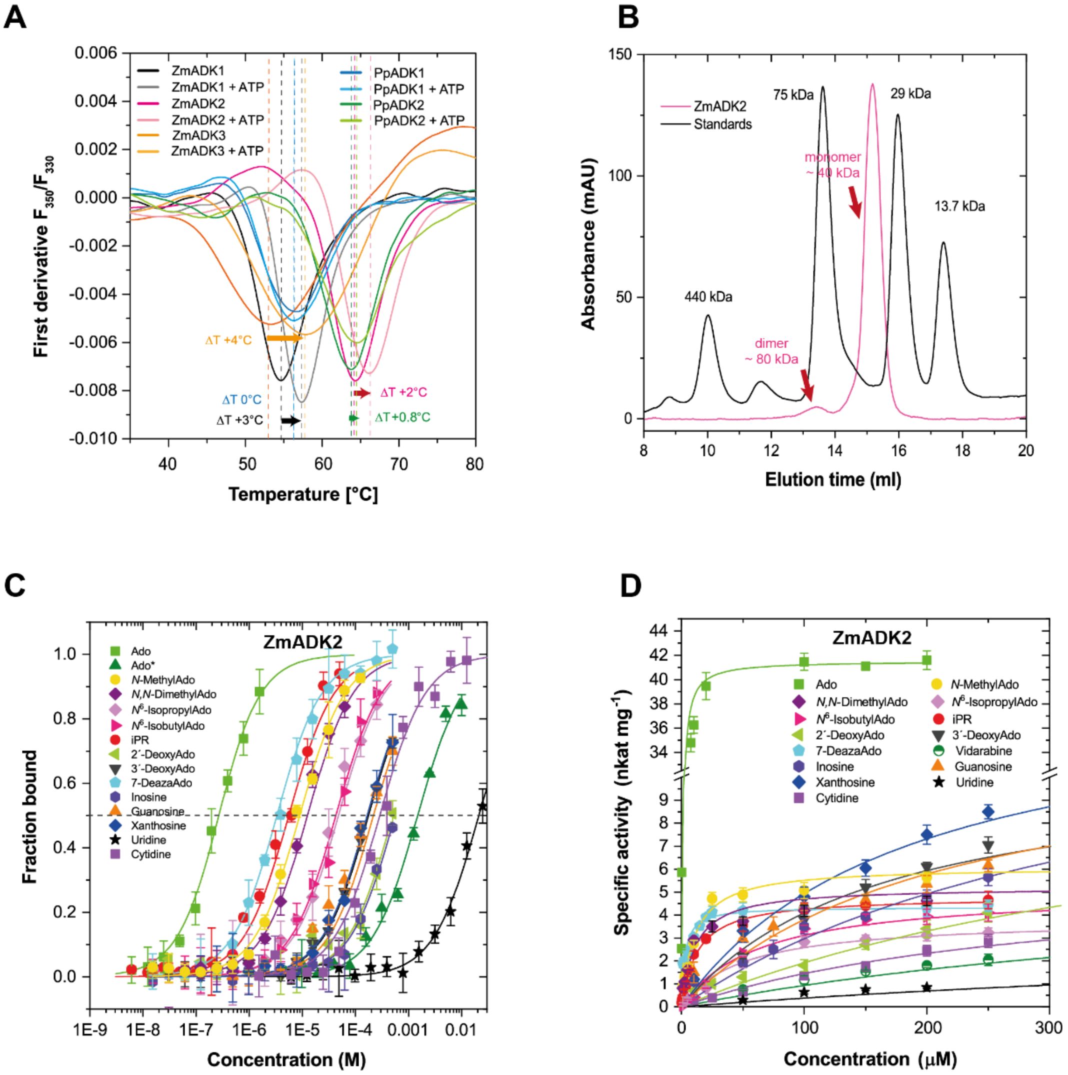
