## Supplementary material for "The monomer/dimer switch modulates the activity of plant adenosine kinase": Figure S2

**Figure S2. Reaction scheme for the synthesis of two purine ribosides.** Purine ribosides were prepared in a one-step reaction by heating 6-chloropurine riboside with the corresponding amine in the presence or absence of triethylamine as an auxiliary base. Reagents and conditions: i) methylamine, ethanol, 90 °C, 4 h; ii) dimethylamine hydrochloride, triethylamine, methanol, 100 °C, 4 h; iii) triethylamine, n-propanol, 85 °C, 4 h and isopropylamine (c) or isobutylamine (d).

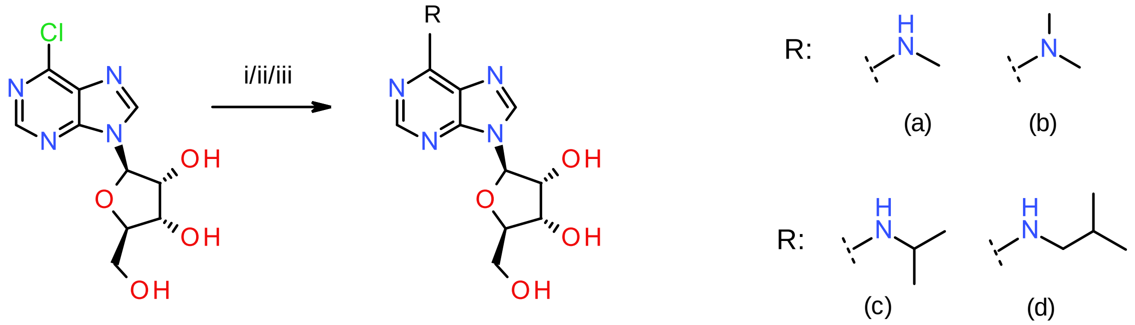


6-chloropurine riboside

*N*[6]-methyladenosine (a)
*N*[6], *N*[6]-dimethyladenosine (b)
*N*[6]-isopropyladenosine (c)
*N*[6]-isobutyladenosine (d)
