## Supplementary figures and images for "The monomer/dimer switch modulates the activity of plant adenosine kinase"

### Figure S3

**Figure S3. NMR spectra of $N^6$-methylAdo.**

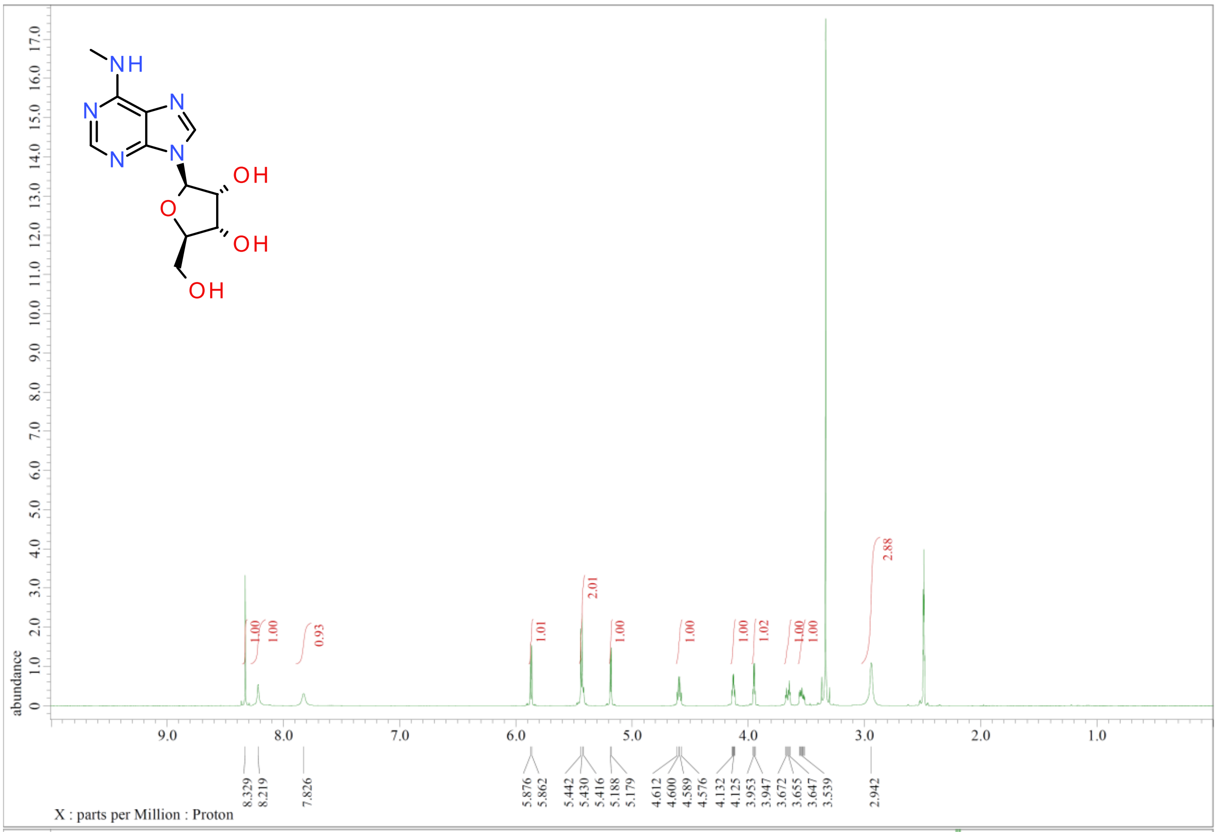


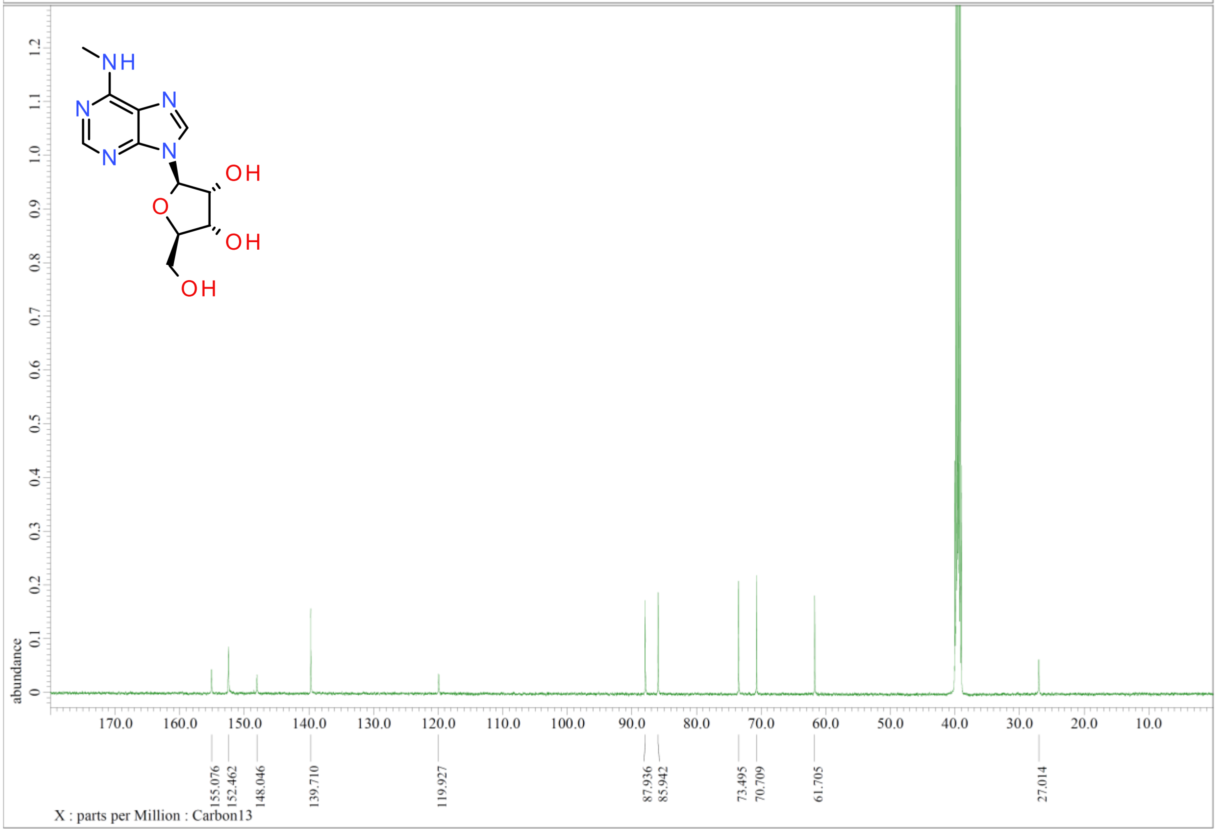

### Figure S4

**Figure S4. NMR spectra of *N*[6], *N*[6]-dimethylAdo.**

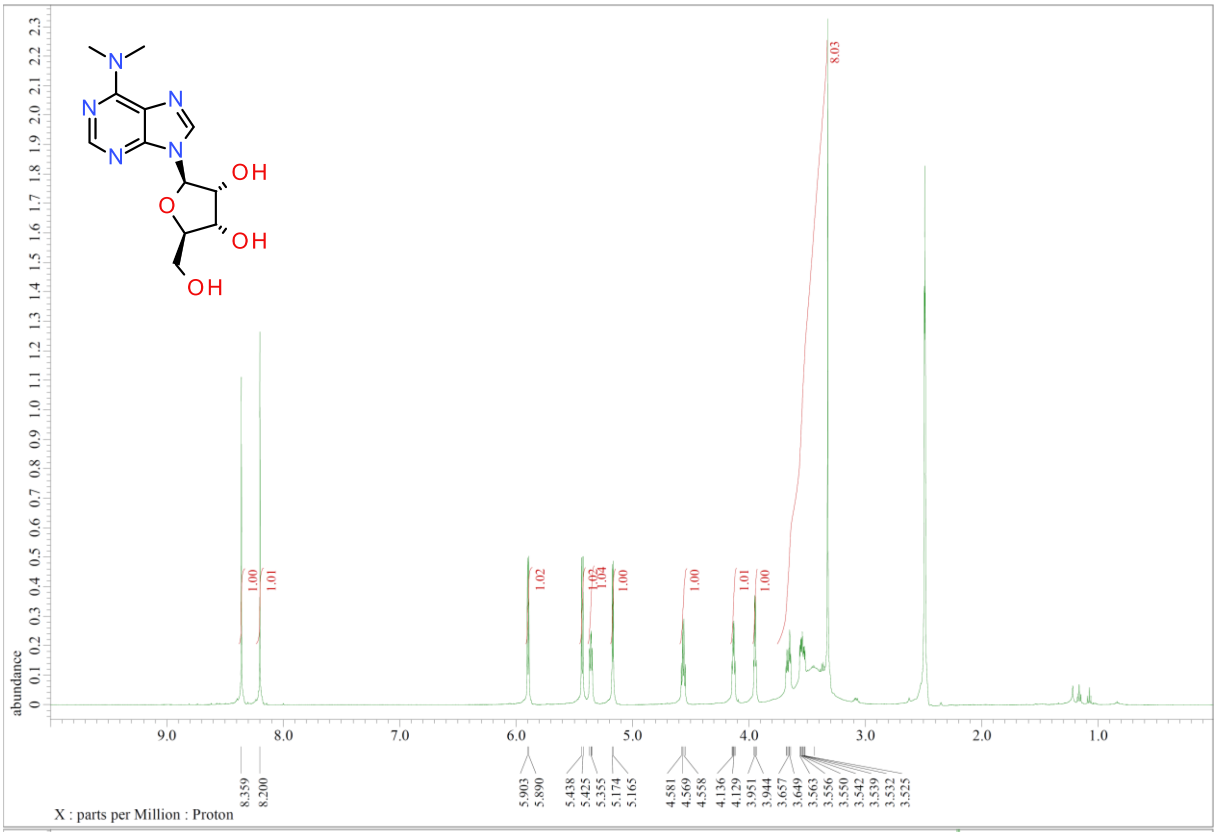


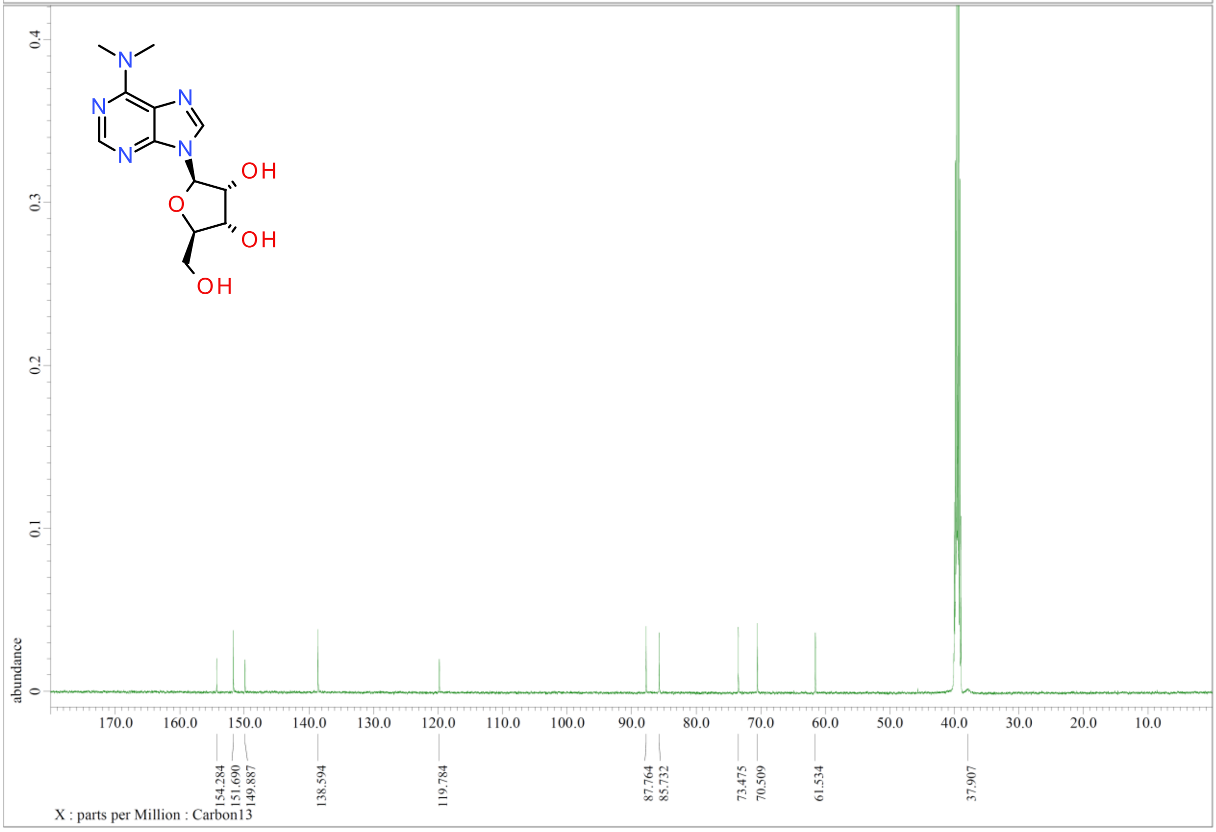

### Figure S5

**Figure S5. NMR spectra of $N^6$-isopropylAdo.**

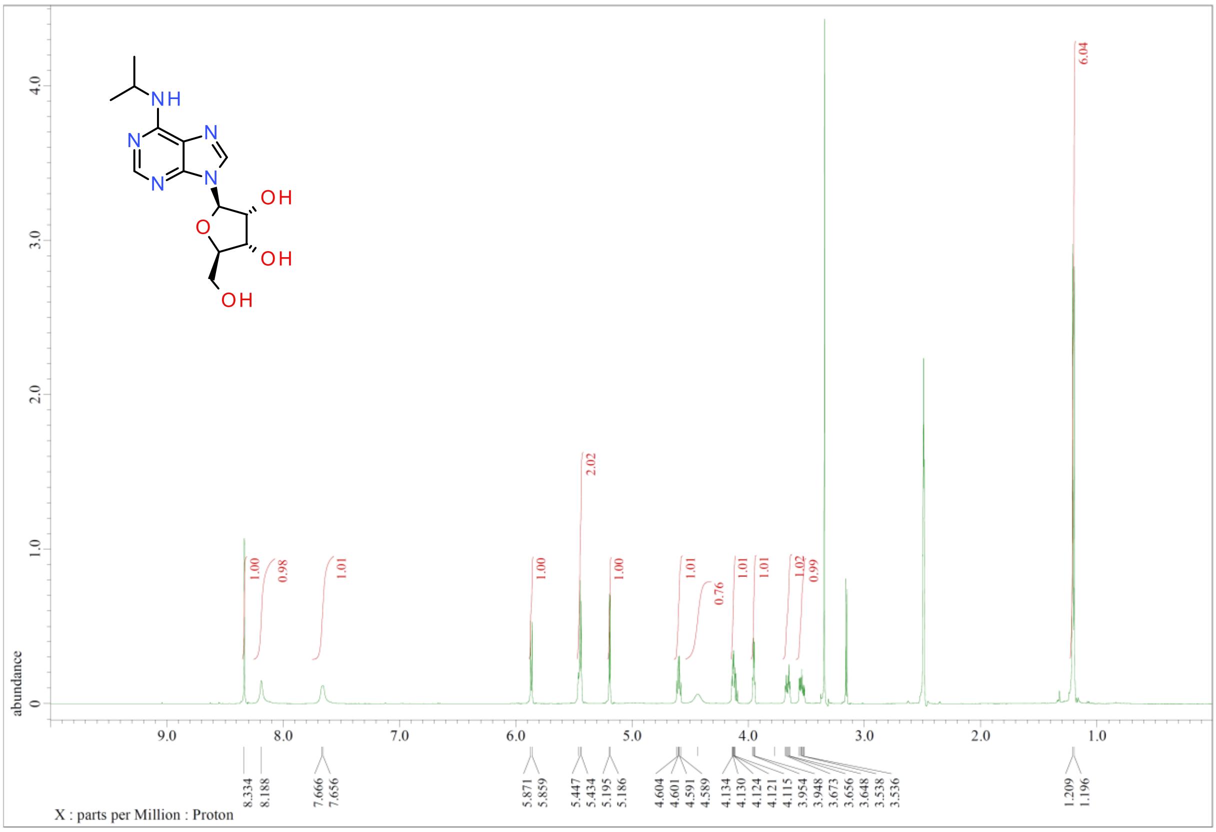


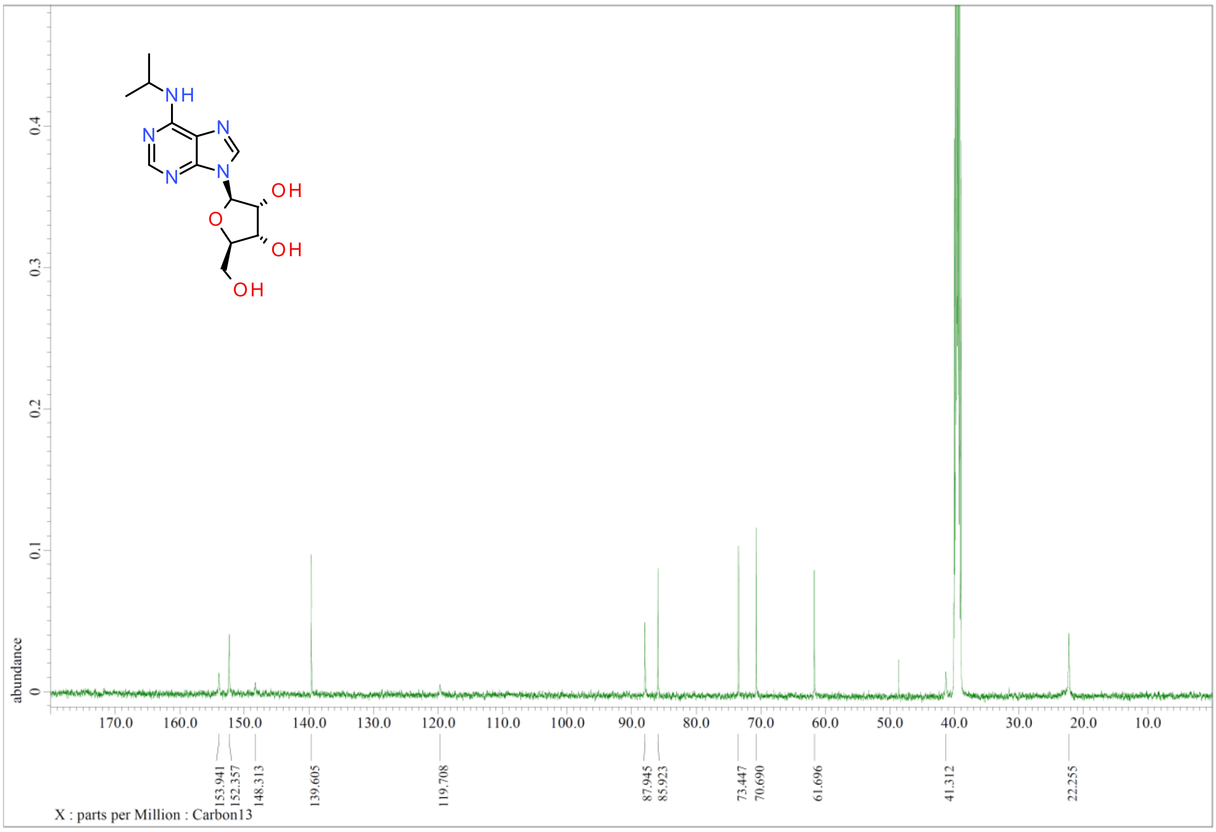

### Figure S6

**Figure S6. NMR spectra of *N*[6]-isobutylAdo.**

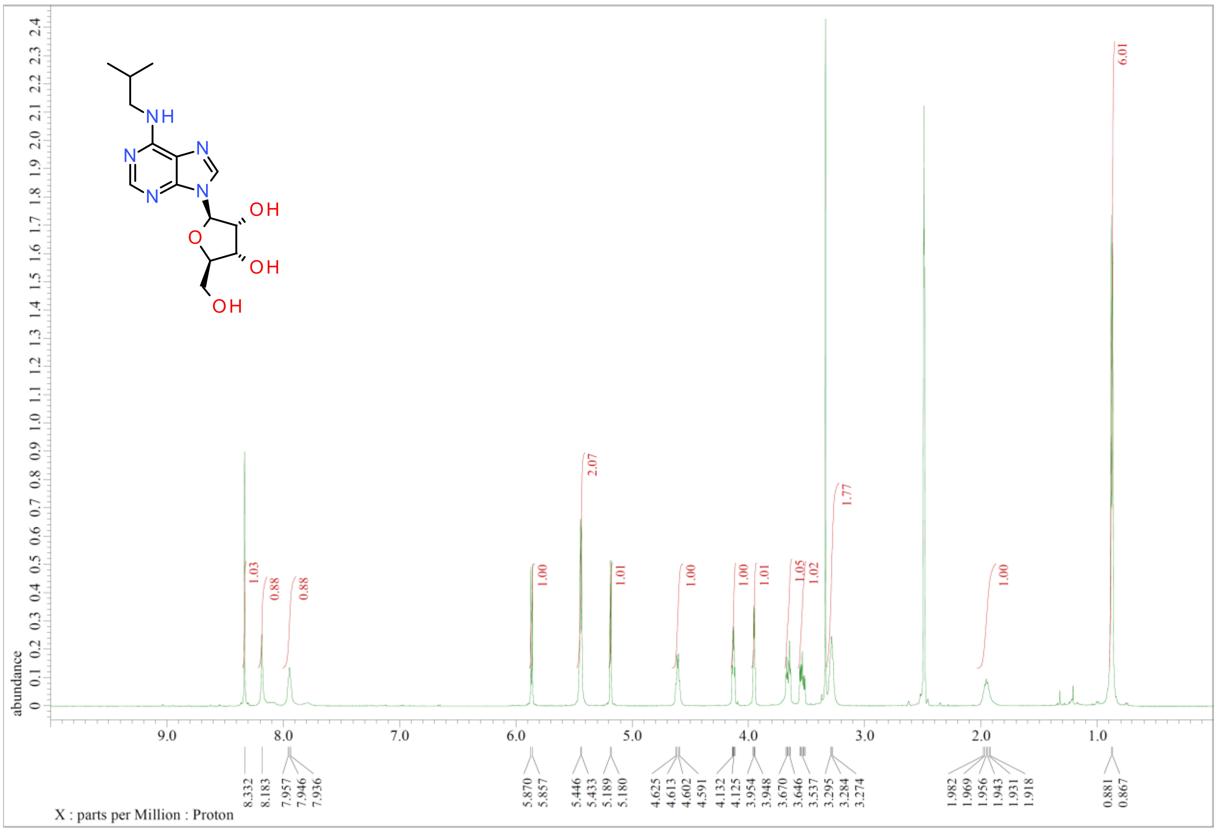


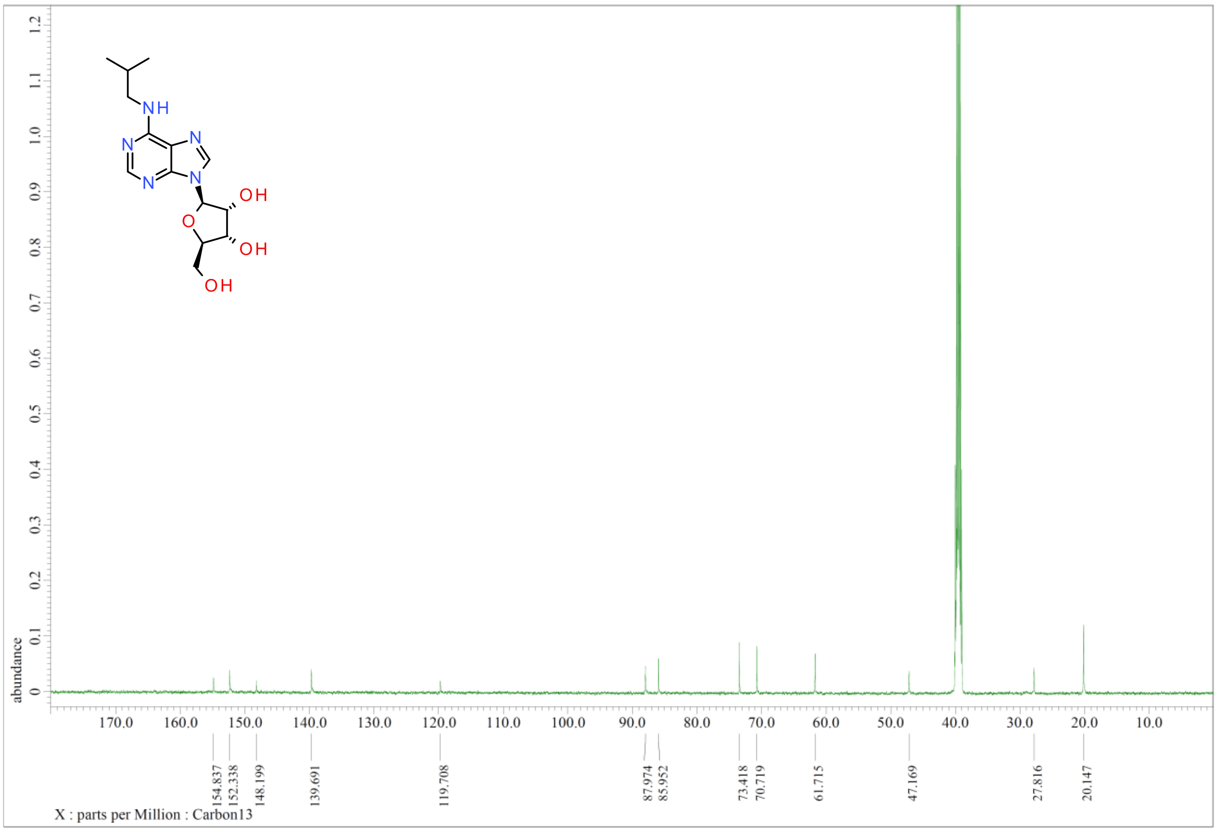
