## Supplementary material for "The monomer/dimer switch modulates the activity of plant adenosine kinase": Figure S7

**Figure S7. Sequence alignment of the selected plant ADKs and human ADK.** Sequence accession numbers are as follows: ZmADK1 (Zm00001d051157), ZmADK2 (Zm00001d017271), ZmADK3 (Zm00001d003017), PpADK1 (Pp3c3_10800), PpADK2 (Pp3c13_10550), PpADK3 (Pp3c8_25260), AtADK1 (At3g09820), AtADK2 (At5g03300) and human ADK (Uniprot accession: P55263). The motif found in ADK sequences is shown in a red rectangle, while residues forming the dimer interface are highlighted in green.

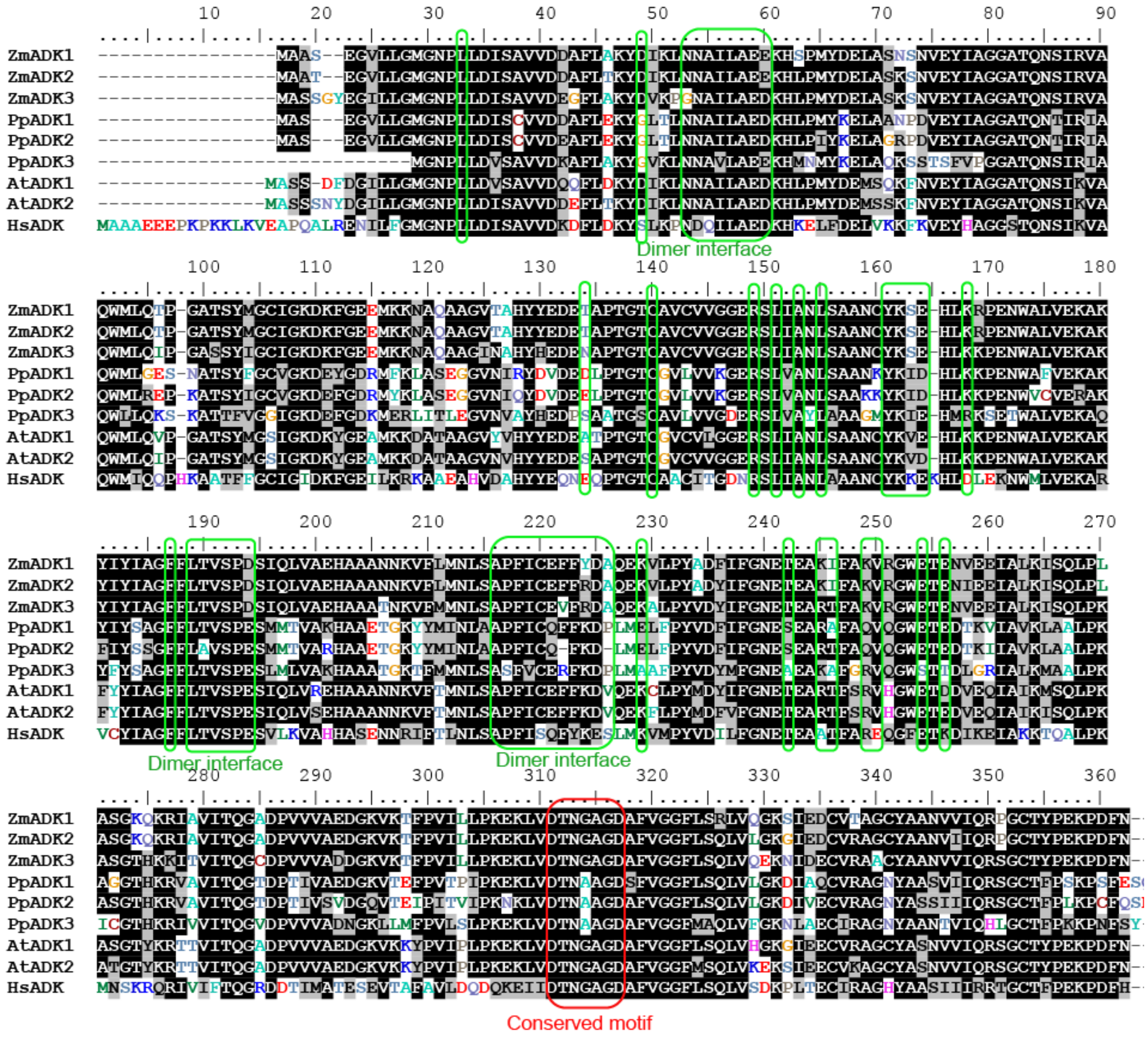
