## Supplementary material for "The monomer/dimer switch modulates the activity of plant adenosine kinase": Figure S8

**Figure S8.** ***In silico docking*** **of cytokinin riboside in the active site of plant ADK**. **(A)** The active site of ZmADK3 (PDB 8RPA) with docked molecules of iPR (green) and Ado (yellow), neighboring residues are labeled, those that were flexible are colored in orange. **(B)** The active site of PpADK1 (PDB 9FW6) with docked molecules of iPR (green) and Ado (yellow). **(C)** The active site of human ADK (PDB 1BX4) with docked molecules of iPR (green) and Ado (yellow), neighboring residues are labeled.

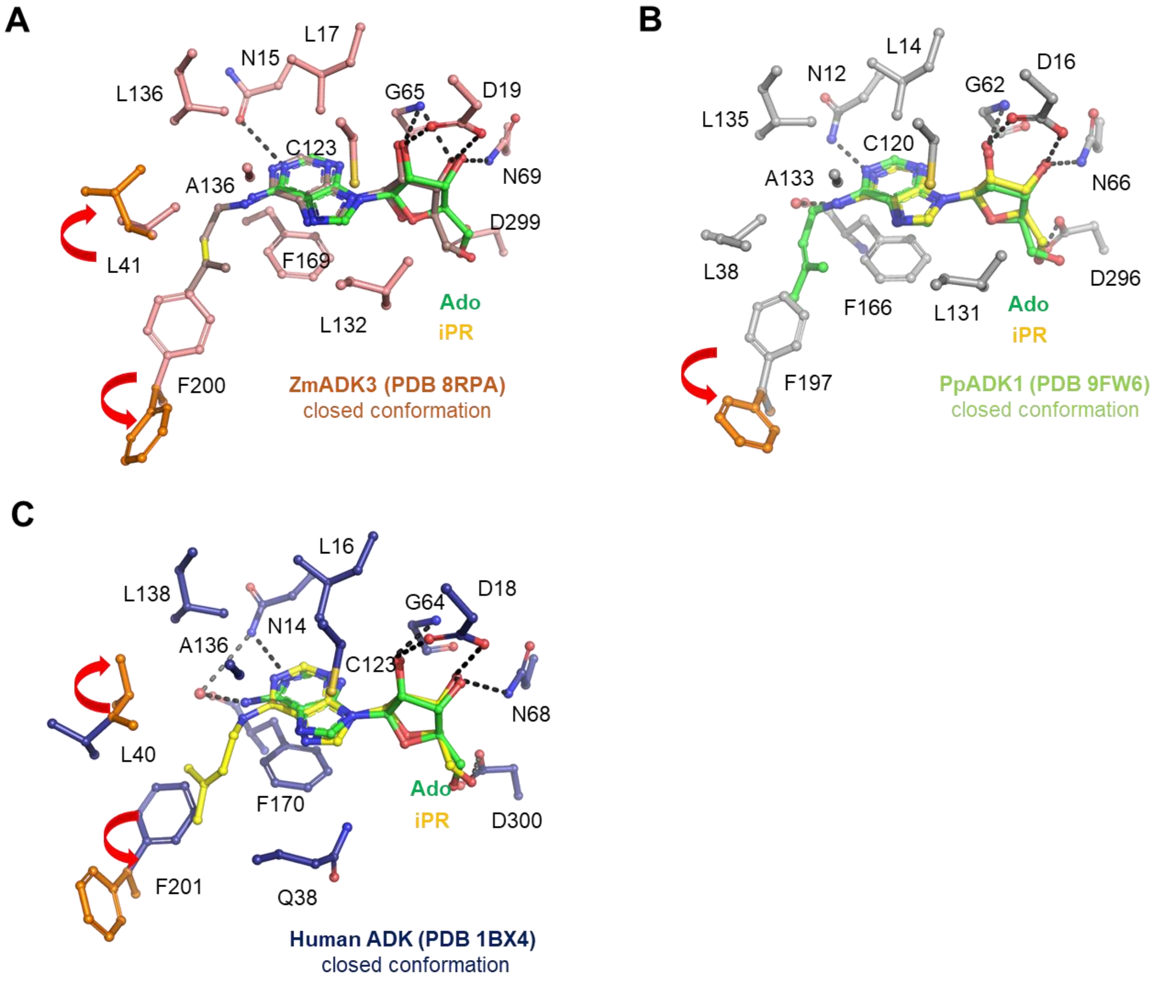
