## Supplementary material for "The monomer/dimer switch modulates the activity of plant adenosine kinase": Figure S9

**Figure S9. Comparison between the two known ADK dimers.** The figure shows **(A)** the surface representation of ADK dimer from *Mycobacterium tuberculosis* in the closed conformation (PDB 2PKK, 22% sequence identity with ZmADKs), **(B)** ADK dimer from *Mycobacterium tuberculosis* in the open conformation (PDB 2PKN) **(C)** plant ADK2 dimer from *Zea mays* in the open conformation (PDB 8RF7, this work).

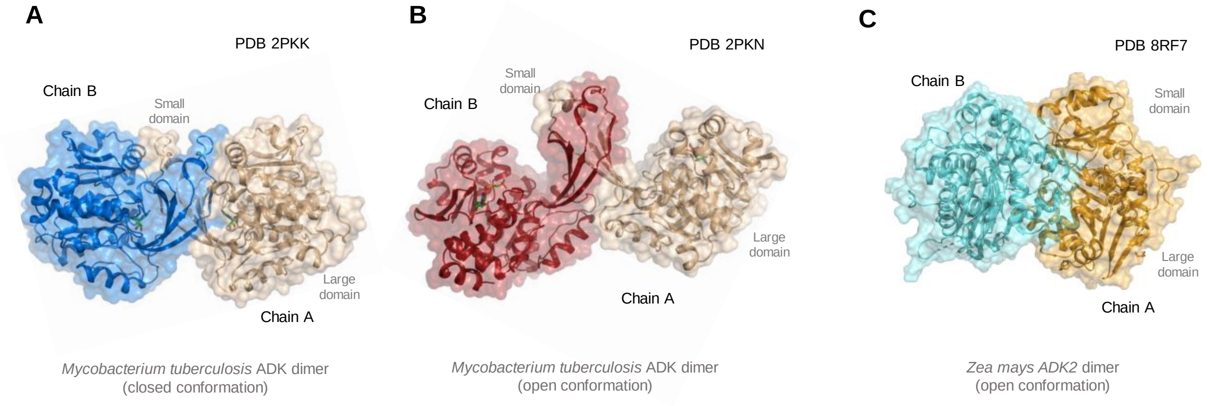
